# Breast tumor stiffness instructs bone metastasis via mechanical memory

**DOI:** 10.1101/847699

**Authors:** Adam W. Watson, Adam D. Grant, Sara S. Parker, Michael W. Harman, Mackenzie R. Roman, Brittany L. Forte, Cody C. Gowan, Raúl Castro-Portuguez, Christian Franck, Darren A. Cusanovich, Megha Padi, Casey E. Romanoski, Ghassan Mouneimne

## Abstract

The mechanical microenvironment of primary breast tumors plays a substantial role in promoting tumor progression^1^. While the transitory response of cancer cells to pathological stiffness in their native microenvironment has been well described^2^, it is still unclear how mechanical stimuli in the primary tumor influence distant, late-stage metastatic phenotypes across time and space *in absentia*. Here, we show that primary tumor stiffness promotes stable, non-genetically heritable phenotypes in breast cancer cells. This “mechanical memory” instructs cancer cells to adopt and maintain increased cytoskeletal dynamics, traction force, and 3D invasion *in vitro*, in addition to promoting osteolytic bone metastasis *in vivo*. Furthermore, we established a mechanical conditioning (MeCo) score comprised of mechanically-regulated genes as a global gene expression measurement of tumor stiffness response. Clinically, we show that a high MeCo score is strongly associated with bone metastasis in patients. Using a discovery approach, we mechanistically traced mechanical memory in part to ERK-mediated mechanotransductive activation of RUNX2, an osteogenic gene bookmarker and bone metastasis driver^3,4^. The combination of these RUNX2 traits permits the stable transactivation of osteolytic target genes that remain upregulated after cancer cells disseminate from their activating microenvironment in order to modify a distant microenvironment. Using genetic, epigenetic, and functional approaches, we were able to simulate, repress, select and extend RUNX2-mediated mechanical memory and alter cancer cell behavior accordingly. In concert with previous studies detailing the influence of *biochemical* properties of the primary tumor stroma on distinct metastatic phenotypes^5–9^, our findings detailing the influence of *biomechanical* properties support a generalized model of cancer progression in which the integrated properties of the primary tumor microenvironment govern the secondary tumor microenvironment, *i.e.*, soil instructs soil.

---

Tumor stiffening is a ubiquitous feature of breast cancer progression which gives rise to a varied mechanical landscape ranging from normal elasticity to fibrotic-like tissue stiffness^10^. When cells engage a stiff matrix through their focal adhesions, the physical resistance triggers mechanotransduction, a rapid conversion of mechanical stimuli into biochemical signals. Mechanotransduction promotes the first steps of metastasis by increasing cytoskeletal dynamics, cell migration, and invasion *in situ*^11^. However, the mechanical stimuli that instruct cell behavior in the primary tumor are not persistent throughout the metastatic cascade, especially when metastatic colonization and outgrowth occur in soft microenvironments (e.g., lung, liver, brain, and bone marrow). Thus, we sought to determine whether cancer cells maintain their mechanically-induced behavior after transition to dissimilar mechanical microenvironments. A similar concept has been established in normal mesenchymal stem cells, which are able to maintain their mechanically-mediated differentiation state after their mechanical microenvironment is altered^12,13^.

First, we screened a panel of breast cancer cell lines and patient-derived xenograft (PDX) primary cells for mechanoresponse in the relevant range for breast tumors^10^, using stiffness-tuned, collagen-coated hydrogels that were either 0.5 kPa (soft) or 8.0 kPa (stiff), and employing well-established mechanosensing and mechanotransduction readouts (differential cell spreading and CTGF gene expression, respectively)^1,14^. While all of the cells tested showed elevated mechanoresponse on stiff hydrogels compared to soft, SUM159 cells showed the greatest combined response (Fig.1b). To interrogate mechanical memory, we developed a multi-functional analysis workflow consisting of a mechanical preconditioning phase and a mechanical memory challenge (or control) phase, followed by multiple functional assays (Fig.1a). First, we examined SUM159 actin cytoskeletal dynamics in three experimental groups: stiff-preconditioned control cells (St7/St1), which are cultured on stiff hydrogels for 7 days in phase 1 (St7) and then transferred to new stiff hydrogels for 1 day in phase 2 (St1); soft-preconditioned control cells (So7/So1); and stiffness-memory cells (St7/So1), which are stiff-preconditioned cells challenged with 1 day on soft hydrogels before analysis. St7/St1 cells exhibited increased cytoskeletal dynamics compared to So7/So1 cells, which maintained reduced dynamics throughout imaging on glass (Fig.1c). Intriguingly, stiffness-memory St7/So1 cells fully retained their increased dynamics and behaved similarly to St7/St1 cells (Fig.1c,d, and Extended Data Fig.1a,b, Supplementary Movie 1). This was recapitulated in the traction-induced displacement signatures obtained via multidimensional traction force microscopy; however, while the magnitude of traction-induced cell displacements was similar among the St7/St1 and St7/So1 cells (Fig.1e and Extended Data Fig.1c,d), the polarization of contractility, or average spatial arrangement of tractions was slightly diminished in both St7/So1 and So7/So1 cells (Fig. 1f and Extended Data Fig.1e). In a live-imaging based invasion assay, stiffness-memory cells retained their stiffness-induced invasive capacity, reflected in the number of invading cells and migration dynamics including speed and directionality (Fig.1g,h and Extended Data Fig.1f-i). Interestingly, although soft-preconditioned control cells invaded less, the translocation of their invasion front was not diminished, suggesting that mechanical memory particularly sustained single-cell dissemination (Fig.1i, Supplementary Movie 2). Together, these results suggest that mechanical memory manifests distinctively in particular cytoskeletal dynamics and is associated with 3D invasion.

**Fig.1.**
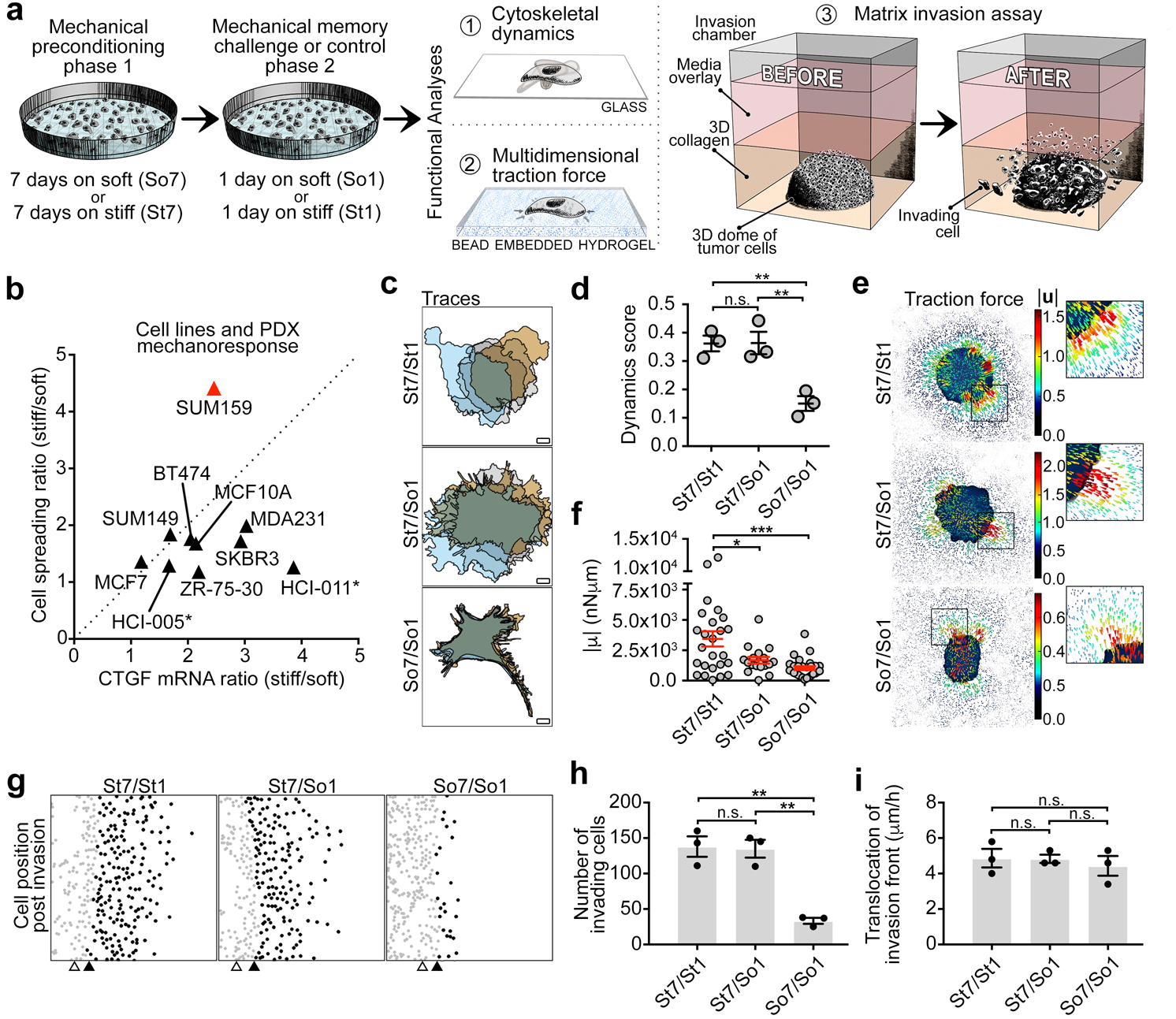
Mechanical memory manifests distinctively in cellular dynamics and invasion. **a**, Schematics showing multifunctional analysis workflow for testing mechanical memory. **b**, Mechanoresponse readouts for a panel of breast cancer cell lines and patient-derived xenografts (denoted by asterisks). **c**, Cytoskeletal dynamics (overlaid cell traces) of iRFP-Lifeact-expressing SUM159 cells preconditioned on stiff and/or soft hydrogels as indicated, showing 1 hour intervals starting 10 hours after plating on glass. Scale bars = 10 µm. See Supplementary Movie 1. **d**, Quantification of (**c**) (*n* = 36 cells in each condition from *n* = 3 biological replicates). **e**, Multidimensional traction force microscopy of SUM159 cells on bead-embedded 8.5 kPa gels, preconditioned on regular stiff and/or soft hydrogels as indicated. Panels show vector maps of displacement magnitude. Heat scales (µm) show bead displacement. **f**, Quantification of (**e**) (*n* = 17-28 cells in each condition from *n* = 3 biological replicates). **g**, SUM159 cell position along invasion fronts after 16 hours of live-cell tracking in 3D collagen. Cells were preconditioned on stiff and/or soft hydrogels as indicated. Gray dots = non-invasive cells; black dots = invasive cells; white triangles = invasion front at start of imaging; black triangles = invasion front at end of imaging. See Supplementary Movie 2, Extended Data Fig.1f. **h**,**i**, Quantification of (**g**) (*n* = 3 biological replicates with *n* = 3 technical replicates). Data are mean ± s.e.m. \**P* < 0.05; \*\**P* < 0.01 \*\*\**P* < 0.001, one-way ANOVA with Tukey’s multiple comparisons test. See Source Data for exact *P* values.

To ascertain the global changes in gene expression in response to fibrotic-like stiffness, we performed differential gene expression analysis on soft- and stiff-preconditioned SUM159 cells using RNA-seq (Fig.2a). Interestingly, examining the pathways upregulated revealed enrichments for a number of skeletal gene ontologies (from Metascape^15^) in the stiffness-induced gene set (Fig.2b and Extended Data Fig.2a). Furthermore, a second pathway enrichment analysis (using Enrichr^16^) revealed an association with numerous skeletal pathologies, through query of the Human Phenotype Ontology and MGI Mammalian Phenotype libraries (Extended Data Fig.2b,c), which led us to consider bone metastasis as a possible phenotypic correlate of stiffness-induced gene expression in metastasis-competent cancer cells.

**Fig.2.**
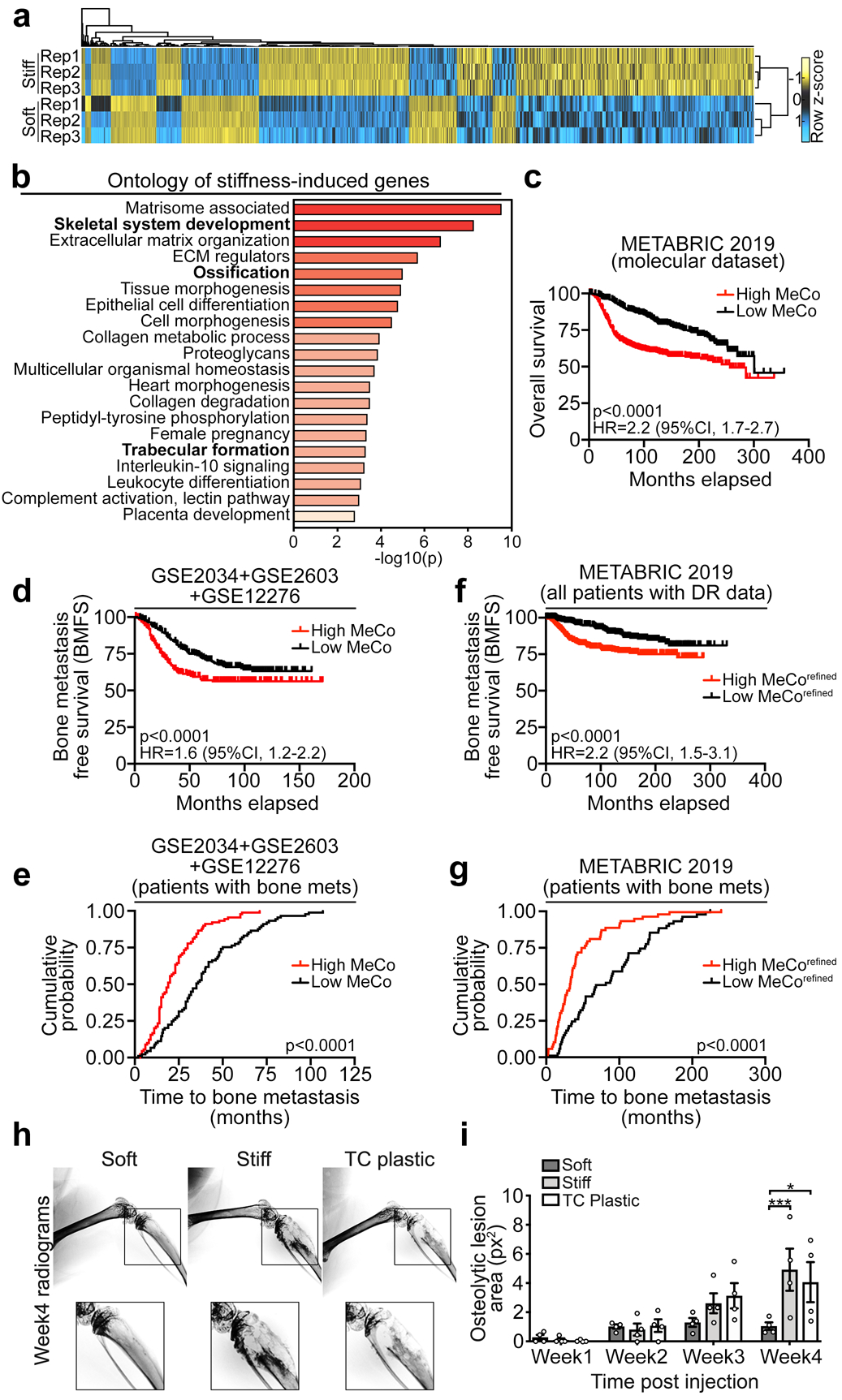
Primary tumor mechanical conditioning is associated with bone metastasis. **a**, Heatmap of differentially-regulated genes (≥2-fold) from RNA-seq of 2-week soft- and stiff-preconditioned SUM159 cells, which constitute the raw MeCo genes (*n* = 3 biological replicates). **b**, Metascape ontology of stiffness-induced genes (>4-fold). Bold text indicates skeletal ontologies. **c**, Kaplan-Meier curve of patients in the METABRIC 2019 study (molecular dataset cohort), assigning each patient a raw “mechanical conditioning” (MeCo) score derived from the differential gene expression in (**a**) comparing upper and lower quartiles of MeCo score (*n* = 476 high MeCo, 476 low MeCo). See Methods for MeCo score derivation. **d**, Kaplan-Meier curve of bone metastasis-free survival in the combined cohort, split at median MeCo score (*n* = 280 high MeCo, 280 low MeCo). **e**, Time to bone metastasis for patients in (**d**) split at median MeCo score (*n* = 93 high MeCo, 92 low MeCo). **f**,**g** Large validation cohort for the MeCo^refined^ score showing Kaplan-Meier curve of bone metastasis-free survival (**f**) and time to bone metastasis (**g**) in METABRIC 2019 (using all patients with distant relapse annotation), comparing upper and lower quartiles of MeCo^refined^ score (*n* = 422 high MeCo^refined^, 421 low MeCo^refined^ in (**f**), and *n* = 65 high MeCo^refined^, 64 low MeCo^refined^ in (**g**). **h**, Radiograms of tibia from mice injected with 7-day soft-, stiff-, and plastic-preconditioned SUM159 cells, imaged 4 weeks after intracardiac injection. **i**, Quantification of (**h**) (*n* = 4 mice per group; TC = tissue culture). Data are mean ± s.e.m \**P* < 0.05; \*\*\**P* < 0.001; \*\*\*\**P* < 0.0001, two-way ANOVA with Tukey’s multiple comparisons test. Kaplan-Meier *P* values calculated with Wilcoxon test. See Source Data for exact *P* values.

To test for a clinical association between breast tumor stiffness and bone metastasis, we developed a mechanical conditioning (MeCo) multi-gene score and used it as a global gene expression measurement of tumor stiffness response. The MeCo score was generated using the entire set of genes that were differentially regulated in response to stiffness in SUM159 cells; however, to mitigate any influence of stiffness-induced proliferation (Extended Data Fig.2d), we excluded genes that are known to be strongly associated with proliferation^17^. We calculated individual MeCo scores for primary breast tumors from multiple patient cohorts. In the large METABRIC 2019 (molecular dataset) cohort, high MeCo scores are strongly associated with poor overall survival (Fig.2c, Hazard Ratio (HR)=2.2, p<0.0001). In a separate, combined cohort of 560 patients for whom metastasis status and site were recorded, high MeCo scores are strongly associated with lower bone metastasis-free survival (BMFS) (Fig.2d, HR=1.6, p<0.0001, Extended Data Fig.2e). Furthermore, amongst the 185 patients in the combined cohort who developed bone metastasis, the median time to bone metastasis (TTBM) was 19 months for those with high MeCo scores, compared to 35 months for those with low MeCo scores (Fig.2e). Interestingly, aggressive PAM50 tumor subtypes, such as basal, HER2 and luminal B, have strikingly higher average MeCo scores compared to the less aggressive luminal A and normal-like subtypes (Extended Data Fig.2f); this is in line with a contrasting expression pattern of PAM50 genes between soft- and stiff-preconditioned cells (Extended Data Fig.2g). These observations are also consistent with our data showing that stiffness-preconditioning enhances invasion. To identify the genes that contribute to the association between MeCo score and bone metastasis independently of subtype and patient cohort, we used linear regression to correct for subtype-specific effects and differences in platform and subtype composition between studies in the combined cohort of 560 patients (Extended Data Fig.3a-c), and we found the subset of MeCo genes (MeCo^refined^) that were consistently up- or down-regulated between bone-metastatic and non-metastatic primary tumors (Extended Data Fig.3d-h). The MeCo^refined^ score was significantly better at predicting BMFS and TTBM than matched gene sets randomly chosen from up- and down-regulated genes between bone-metastatic and non-metastatic primary tumors (Extended Data Fig.3g,h). More importantly, the MeCo^refined^ score was validated in two independent datasets, NKI (HR=2.1, p<0.009) and METABRIC 2019 (HR=2.2, p<0.0001) (Fig.2f,g and Extended Data Fig.3i,j). In the latter, the median TTBM for patients with high MeCo^refined^ scores was 33 months, compared to 85 months for those with low MeCo^refined^ scores (Fig.2g). Together, these analyses reveal how mechanical conditioning is associated with bone metastasis at the genome-wide level.

To determine if fibrotic-like stiffness promotes bone metastasis *in vivo*, we injected NOD- *scid IL2rγ*^*null*^ mice with soft-, stiff- or plastic-preconditioned SUM159 cells in the left cardiac ventricle and tracked bone changes with X-ray imaging. Progressive osteolysis was evident in both the stiff- and plastic-preconditioned groups compared to the soft-preconditioned group (Fig.2h,i, Extended Data Fig.4a). To test the osteolytic capacity of stiffness-memory cells, and to simultaneously determine whether the colonization deficit in the soft-preconditioned group was extravasation-dependent, we performed intrafemoral injection of St7/So1 or So7/So1 cells in order to bypass systemic circulation. Stiffness-memory St7/So1 cells grew faster *in situ*, and their growth led to reduced cortical bone volume (Extended Data Fig.4b-g). These data strongly support that mechanical memory promotes osteolytic disease *in vivo*.

We undertook a function-based discovery approach to identify candidate drivers of mechanical memory-regulated bone metastasis. Focusing on the induction of transcriptional memory by mechanotransduction, we identified upstream transcriptional regulators using Ingenuity Pathway Analysis and the mechanically-induced gene set. Next, we cross-listed these regulators with a large database of metastasis-associated genes^18^; within the list of common genes, we selected those with evidence of transcriptional memory function, with the rationale that such candidates would offer the type of phenotype durability we observe after mechanical conditioning. This yielded 5 candidates (Fig.3a). Among these, the osteogenic transcription factor RUNX2 was especially of interest because of its well-described roles in invasion, bone metastasis and genomic bookmarking, which selectively maintains open chromatin signatures during cell division, thus promoting transcriptional memory in daughter cells^3,4,19^. Using the assay for transposase-accessible chromatin, ATAC-seq^20^, we examined the dynamics of chromatin accessibility following the transition from stiff to soft matrices over a 7-day time course, *i.e.*, loss of mechanotransduction (Extended Data Fig.5a). As expected, the similarity in epigenomes across samples was largely driven by the amount of time cells were conditioned in a soft environment (Fig.3b and Extended Data Fig.5b,c). To better understand the epigenetic code underlying dynamic differences during environmental conditioning, we focused on two sets of accessible sites that close upon transition to soft substrate, yet with different patterns of change: a ‘delayed-closing’ set, and a ‘quick-closing’ set. Using a likelihood ratio test framework, delayed-closing sites were considered as those with no significant change in accessibility by day 2 on soft matrix, but with significantly less accessibility by day 5. In contrast, quick-closing sites were those exhibiting a significant loss of accessibility after only 12 hours in the soft environment, which is a period of time shorter than the mean population doubling (Extended Data Fig.5d). We identified 7,277 delayed-closing genomic regions, which we reasoned should be enriched for regulatory sites responsible for mechanical memory. Using a genomic region-based pathway enrichment analysis^21^, we found that these sites are implicated in ‘abnormal bone remodeling’ (q=0.015), among other pathways (Supplementary Information Table 5). Furthermore, using unbiased motif enrichment analysis for the sequences at these loci, we identified that the RUNX consensus binding motif is enriched in delayed-closing (p=1e-33), but not quick-closing sites (Fig.3c); importantly, RUNX binding motifs were also significantly enriched in sites that maintained their accessibility throughout our time course after transitioning to soft (p=1e-1458) (Supplementary Information Table 6-8). Together, these data are consistent with the bookmarking function of RUNX2 and its role in promoting mechanical memory. By comparison, the consensus binding motif of BACH (known regulator of bone metastasis^22^) is enriched in the quick-closing sites, suggesting that it does not play a role in mechanical memory. These analyses prioritized RUNX2 for further investigation.

**Fig.3.**
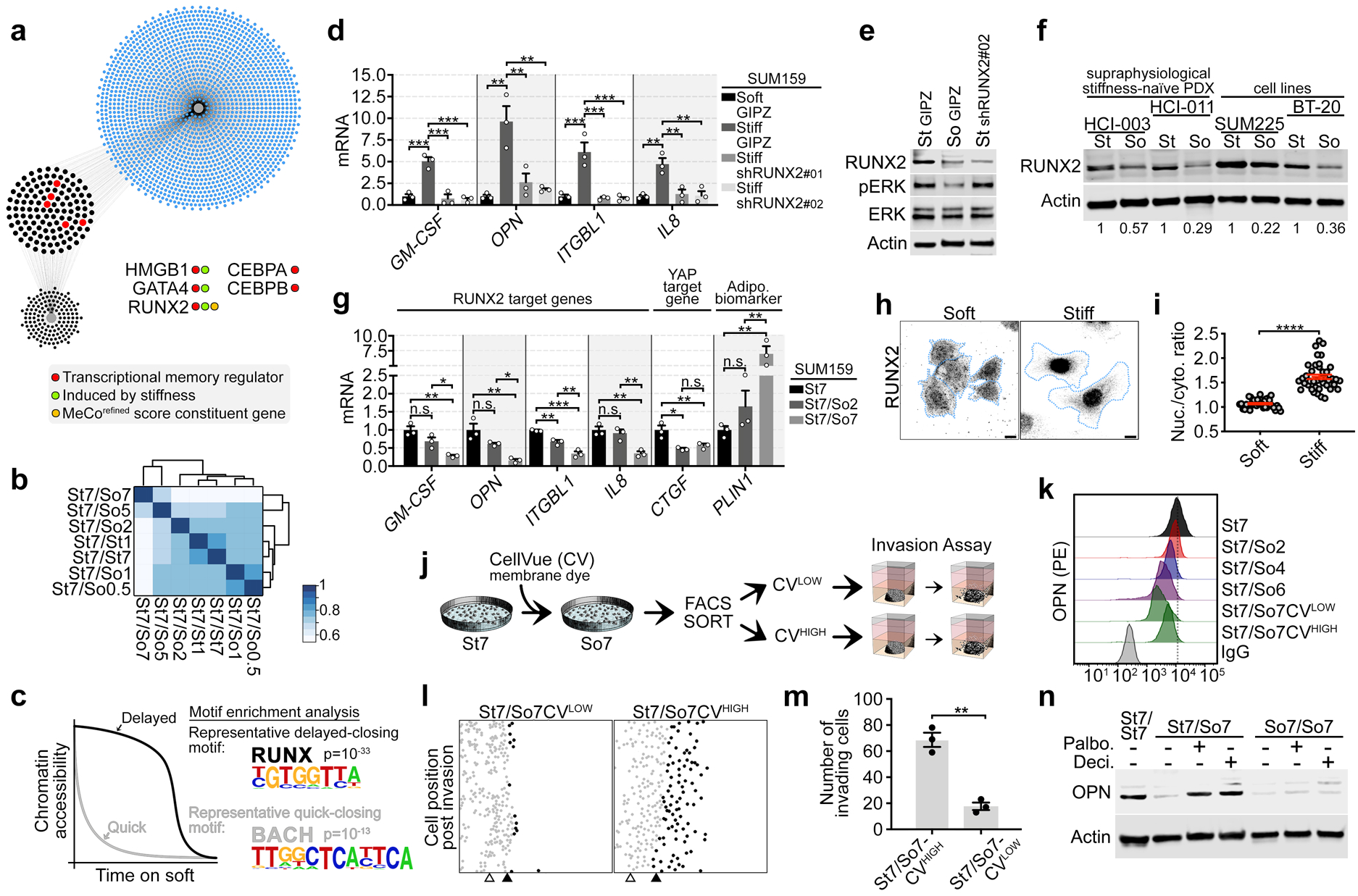
RUNX2 is activated by stiffness and is associated with proliferation-sensitive mechanical memory. **a**, Schematics of discovery approach identifying candidate drivers of mechanical memory-mediated metastasis. Gray dots = mechanically-sensitive upstream regulators from RNA-seq analysis (141 genes); blue dots = Human Cancer Metastasis Database metastasis-associated genes (1811 genes); black/red dots = intersecting genes (123 genes). Red dots = intersecting genes that are known gene bookmarkers (5 genes) and inset shows further characterization. See Methods for detailed description. **b**, Heat map showing clustering of ATAC-seq samples; scale shows Jaccard index. See Extended Data Fig.5a for sample annotation. **c**, Graphic representation of change in chromatin accessibility over time after changing the mechanical environment; inset shows RUNX and BACH family motifs as representative of the significantly enriched motifs in the delayed-closing and quick-closing sites, respectively (motifs analysis performed by HOMER). **d**, RT-qPCR of 4 RUNX2 target genes in SUM159 cells preconditioned for 7 days on soft and stiff hydrogels with non-targeting shRNA (GIPZ), or on stiff hydrogels with two shRNAs targeting RUNX2 (*n* = 3 biological replicates). Data are mean ± s.e.m. \**P* < 0.05; \*\**P* < 0.01; \*\*\**P* < 0.001, one-way ANOVA with Tukey’s multiple comparisons test. **e**, Immunoblot of RUNX2, ERK and pERK in SUM159 cells stably expressing lentiviral shRUNX2 or GIPZ (non-targeting control), preconditioned for 7 days on soft or stiff hydrogels (representative of *n* = 3 biological replicates). **f**, Immunoblot of RUNX2 in patient-derived xenograft (PDX) primary cells and breast cancer cell lines, preconditioned on soft and stiff hydrogels for 7 days (representative of *n* = 2 biological replicates). **g**, RT-qPCR of RUNX2 and 4 target genes, plus CTGF (YAP target) and PLIN1 (adipogenic biomarker) in SUM159 cells preconditioned as indicated (*n* = 3 biological replicates). Data are mean ± s.e.m. \**P* < 0.05; \*\**P* < 0.01; \*\*\**P* < 0.001, one-way ANOVA with Tukey’s multiple comparisons test. **h**, Immunofluorescence staining of RUNX2 in SUM159 cells on soft and stiff hydrogels. Cell boundaries are delineated in blue. Scale bars = 10 µm. **i**, Quantification of (**e**) (*n* = 40 cells in each condition from *n* = 3 biological replicates). ****P < 0.0001, two-tailed unpaired Student’s *t*-test. **j**, Schematics showing strategy to enrich high-proliferative cells (CV^LOW^) vs low-proliferative cells (CV^HIGH^). **k**, Time course of mechanical memory loss, showing flow cytometry of SUM159 cells preconditioned as indicated, sorted as in (**g**) and stained for OPN (*n* = 3 biological replicates). See Extended Data Fig.8. **l**,**m**, SUM159 cell position (**i**) and quantification (**j**) after 16 hours of live-cell tracking in 3D collagen. Gray dots = non-invasive cells; black dots = invasive cells; white triangles = invasion front at start of imaging; black triangles = invasion front at end of imaging. See Supplementary Movie 3, Extended Data Fig.8b,c. Cells were preconditioned as outlined in (**g**) (*n* = 3 biological replicates with *n* = 3 technical replicates). Data are mean ± s.e.m. \*\**P* < 0.01, two-tailed unpaired Student’s *t*-test. **n**, Immunoblot of OPN showing memory extension in SUM159 cells treated with DMSO, palbociclib (2.5 µM; CDK4/6 inhibitor) or decitabine (7 µM; DNMT1 inhibitor) on soft hydrogels for 7 days in phase 2, after stiff- or soft-preconditioning for 7 days in phase 1. Drugs were added only in phase 2 (representative *n* = 3 biological replicates). See Source Data for exact *P* value.

In a panel of cell lines and PDX primary cells we observed increased RUNX2 in stiff- vs soft-preconditioned cells, in both 2D and 3D culture systems (Fig.3e,f and Extended Data Fig.5f,g). We next measured expression of known RUNX2 target genes^23–25^, and found that the osteolytic genes GM-CSF, OPN, ITGBL1 and IL8, were induced by stiffness in a RUNX2-dependent manner (Fig.3d and Extended Data Fig.5e). Consistent with mechanical memory, upon removal from stiffness, RUNX2 target transactivation was only gradually lost, concomitant with a gradual increase in PLIN1, an adipogenic biomarker (Fig.3g). YAP target expression did not persist similarly to RUNX2 targets (Extended Data Fig.9k), which was surprising since YAP has been linked to mechanical memory in epithelial cells^26^. However, despite being a well-established transducer of mechanical cues, YAP has no known gene bookmarking function^19,27^.

Mechanistically, we observed increased ERK activation on stiff hydrogels, and we confirmed that expression of RUNX2 targets is downstream of stiffness-induced ERK activation of RUNX2; moreover, this was dependent on the activity of the classic mechanotransduction mediators, Src and FAK, in addition to the contractile activity of the actomyosin cytoskeleton (Extended Data Fig.5h-l). On the other hand, inhibition of AKT, another regulator of RUNX2^28^, did not similarly suppress RUNX2 targets at 8 kPa stiffness (Extended Data Fig.5m,n).

To further validate the involvement of RUNX2 in mechanical memory, we generated MCF10A-Neu-RUNX2 cells by expressing human RUNX2 in *Neu*-transformed MCF10A cells^29^, which express low levels of endogenous RUNX2 (Extended Data Fig.10d). The expression of the osteolytic RUNX2 target genes was induced by exogenous RUNX2 on a stiff substrate and remained significantly elevated after two days on soft, compared to seven days on soft (Extended Data Fig.5o). These data are consistent with the role of RUNX2 in promoting mechanical-memory.

With respect to localization, RUNX2 was retained in the cytoplasm in both soft-preconditioned cells on glass and soft-cultured cells *in situ*, compared to stiff-preconditioned cells on glass and stiff-cultured cells *in situ*. This localization pattern was independent of cell spreading, suggesting that it is not due to volume effect (Fig.3h,i, Extended Data Fig.5p, 6a-h). Importantly, in supraphysiological stiffness-naïve HCI-005 PDX primary cells, nuclear RUNX2 was increased on stiff hydrogels compared to soft, demonstrating that this phenotype is not restricted to plastic-tolerant cell lines (Extended Data Fig.6c,d). Lastly, previous studies have demonstrated that RUNX2 is retained in the cytoplasm by stabilized microtubules^30^; by pharmacologically stabilizing actin or tubulin in stiff-cultured cells, or destabilizing the cytoskeleton in soft-cultured cells, we were able to confirm this finding through the expected directional changes in cytoplasmic retention of RUNX2 (Extended Data Fig.7a-d).

As a bookmarking transcription factor, RUNX2 remains bound to chromatin through mitosis to maintain cellular phenotype across cell generations^3^. However, without continual activation by matrix stiffness, we reasoned that RUNX2 activity may decrease after multiple cell divisions. Therefore, we hypothesized that mechanical memory might be erased via proliferation. To test this, we preconditioned cells on stiff hydrogels for 7 days to encode mechanical memory, and then labelled them with CellVue (a membrane dye retention approach to track proliferation) just before switching them to soft hydrogels for 7 days (Fig.3j, Extended Data Fig.8a). RUNX2 target expression was higher in the low-proliferative St7/So7Cellvue^HIGH^ cells compared to St7/So7Cellvue^LOW^ cells (Fig.3k, Extended Data Fig.8d,e). In addition, St7/So7Cellvue^HIGH^ cells retained greater invasive ability than St7/So7Cellvue^LOW^ cells (Fig.3l,m, Extended Data Fig.8b,c and Supplementary Movie 3). To determine if long-term memory is causally related to reduced proliferation, we pharmacologically reduced cell-cycle progression directly and indirectly. Inhibition of CDK4/6 extended OPN expression upon removal from stiffness, yet it did not induce expression *de novo* in soft-preconditioned cells that had already lost their memory (Fig.3n). Similar results were obtained when we inhibited DNMT1 (Fig.3n), which is also consistent with reports showing OPN expression is regulated by promoter methylation^31^. Together, these results suggest that in mechanically-sensitive cells, RUNX2 activity is associated with the maintenance of mechanical memory, and low-proliferative cells possess more durable mechanical memory.

In contrast to SUM159 cells, SKBR3 breast cancer cells did not exhibit substantial changes in cytoskeletal dynamics or invasion in response to mechanical conditioning (Extended Data Fig.9a-g). This correlates with their limited tumorigenicity and low metastatic potential *in vivo*^32^. We derived a new subline, MS-SKBR3.1 cells, by extended culture in a mechanical sensitization media containing ascorbic acid and phosphate, which are osteogenic factors known to be elevated in breast tumors^33,34^. In MS-SKBR3.1 cells, stiffness induced RUNX2 localization to the nucleus, and it triggered mechanical memory that was reflected in cytoskeletal dynamics, RUNX2 target expression, and 3D invasion (Extended Data Fig.9a-j, Supplementary Movie 4,5). Together, these data further illustrate the link between pathological stiffness response, RUNX2 activity, and bone metastatic competency.

To determine the role of RUNX2-mediated mechanical memory in metastasis, we generated RUNX2 point mutants at two crucial ERK phosphorylation sites: RUNX2-S301A-S319A (RUNX2-SA), which is a non-phosphorylatable mutant that is unresponsive to ERK stimulation, and RUNX2-S301E-S319E (RUNX2-SE), which is a phosphomimetic mutant that exhibits high transcriptional activity irrespective of ERK stimulation^35^. Crucially, phosphorylation of these sites by ERK is necessary for the epigenetic modification of chromatin by RUNX2^36^, central to its role in transcriptional memory^37^. RUNX2-WT cells had higher RUNX2 target expression on stiff hydrogels compared to soft, indicating that stiffness can activate overexpressed wild-type RUNX2; in addition, RUNX2-SE cells on soft hydrogels had partially rescued stiffness-induced target expression, while RUNX2-SA cells on stiff hydrogels had dramatically reduced target expression (Fig.4a, Extended Data Fig.10c-e). These changes in RUNX2 activity were reflected in changes in invasiveness *in vitro* and in osteolytic bone metastasis after intracardiac injections *in vivo* (Fig.4b-d, Supplementary Movie 6, Extended Data Fig.10a-c, 11a-e). Notably, non-osseous metastases, *i.e.*, lung, liver and brain, showed dissimilar patterns of disease burden (Extended Data Fig.11f-j). In addition, since RUNX2 targets (particularly OPN) are known to mediate adhesion to bone matrix, and to activate osteoclasts^4,38,39^, we sought to determine how stiffness-activated RUNX2 affected these parameters. Compared to soft-preconditioned, stiff-preconditioned RUNX2-WT cells adhered/spread faster and protruded more on synthetic bone matrix, and induced more paracrine osteoclastogenesis (Fig.4e, Supplementary Movie 7, Extended Data Fig.11k-p). This phenotype was mimicked in soft-preconditioned RUNX2-SE cells, and repressed in stiff-preconditioned RUNX2-SA cells, confirming the importance of stiffness-induced phosphorylation in promoting mechanical memory.

**Fig.4.**
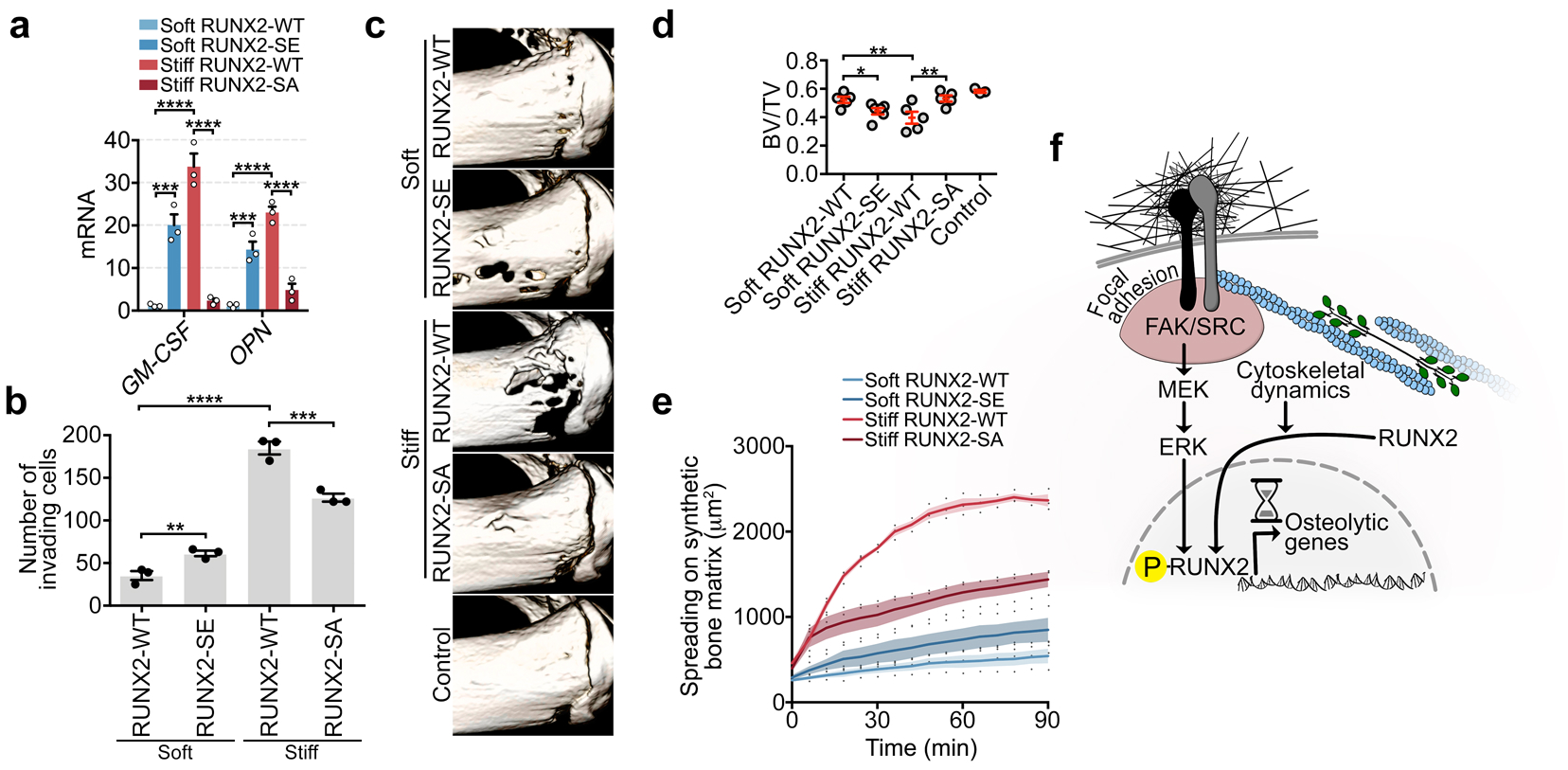
RUNX2-mediated mechanical memory instructs bone metastasis. **a**, RT-qPCR of RUNX2 target genes in SUM159 cells overexpressing RUNX2-WT, RUNX2-SE, or RUNX2-SA, preconditioned for 7 days on soft or stiff hydrogels (*n* = 3 biological replicates). **b**, Quantification of invasion of SUM159 cells preconditioned as indicated in (**a**). (*n* = 3 biological replicates with *n* = 3 technical replicates). See Supplementary Movie 6, Extended Data Fig.10a,b. **c**, Micro-CT 3D reconstructions of proximal tibia from mice 4 weeks after intracardiac injection of SUM159 cells preconditioned as in (**a**) or no cancer cells (control). **d**, Micro-CT analysis of bone volume from mice in (**c**) (*n*: mice; soft RUNX2-WT 5; soft RUNX2-SE 6; stiff RUNX2-WT 5; stiff RUNX2-SA 5; control 3). **e**, Time course of SUM159 cells spreading on synthetic bone matrix, preconditioned as in (**a**) (*n* = 36 cells in each condition from *n* = 3 biological replicates). See Supplementary Movie 7. Shaded regions are mean ± s.e.m. \*\*\*\**P* < 0.0001, two-way ANOVA with Tukey’s multiple comparisons test **f**, Model of stiffness-induced RUNX2 activation. Data are mean ± s.e.m. \**P* < 0.05; \*\**P* < 0.01; \*\*\**P* < 0.001; \*\*\*\**P* < 0.0001, one-way ANOVA with Holm-Sidak’s multiple comparisons test except for (**e**). See Source Data for exact *P* values.

In our working model (Fig.4f), mechanical memory is mediated by RUNX2, a mechanically-sensitive gene bookmarker. RUNX2 is activated by fibrotic-like stiffness, first by nuclear localization following increased cytoskeletal dynamics, and then by mechanotransduction via ERK phosphorylation. Mechanical memory can be mimicked in soft-preconditioned cells with overexpression of a phosphomimetic RUNX2 mutant, and repressed in stiff-preconditioned cells with overexpression of a nonphosphorylatable RUNX2 mutant. RUNX2-mediated mechanical memory is necessary and sufficient to enhance adhesion to synthetic bone matrix, to activate osteoclasts, and to promote osteolytic bone metastasis. Mechanical memory is lost gradually upon removal from the encoding microenvironment, concomitant with proliferation. A corollary to this relationship between proliferation and mechanical memory is that osteolytic capacity upon exit from dormancy may be a latent function of this phenomenon. Furthermore, since cancer-induced osteolysis is perpetuated by a positive-feedback loop between osteoclasts and cancer cells known as the “vicious cycle,” a crucial role for mechanical memory may be to kick-start this process^40^.

This study demonstrates that mechanical conditioning is associated with bone metastasis in animal models and clinical data. Since bone metastases inflict the greatest morbidity associated with breast cancer and become incurable in a majority of women with advanced disease^41^, it is critical that we strive to predict their genesis. The MeCo score we present here is a proxy for assessing tumor stiffness *response*, rather than stiffness itself. This distinction is advantageous since most invasive breast tumors are stiffer than surrounding tissue^42^, yet only a fraction produce osteolytic metastases.

