## Extended Data for "Breast tumor stiffness instructs bone metastasis via mechanical memory"

### METHODS

#### CELL CULTURE

SUM149 and SUM159 cells were cultured in Ham's F12 media (Corning) supplemented with 5% HI-FBS (Gibco), 5 μg/ml insulin (Roche), 1 μg/ml hydrocortisone (Sigma) and antibiotics (100 units/mL penicillin + 100 μg/ml streptomycin from Life Technologies, Inc.). MDA-MB-231, HEK293T, BT20, T47D, ZR-75-30, MCF7, BT474 and PC3 cells were cultured in DMEM high glucose media (Corning) supplemented with 10% FBS and antibiotics. HCI-005, HCI-011, HCI-003 and were cultured in Mammocult base media (Stemcell Technologies) supplemented with Mammocult additives. MCF10A cells were cultured in DMEM/F12 media (Corning) with 5% horse serum, 20 ng/ml EGF, 0.5 mg/ml hydrocortisone, 100 ng/ml cholera toxin, 10 μg/ml insulin plus antibiotics. SKBR3 cells were cultured in McCoy's 5A modified media (Corning) supplemented with 10% FBS and antibiotics. MCF10A-Neu cells were a gift from Cheuk Leung (University of Minnesota). MS-SKBR3.1 cells were derived from SKBR3 cells cultured for 6 weeks at 80-100% confluency in DMEM high glucose media supplemented with 10% FBS, 50 μM ascorbic acid 2-phosphate, and 5 mM disodium glycerol-2-phosphate plus antibiotics (mechanical sensitization media; M.S. media). For live-cell imaging experiments, cells were maintained in a climate-controlled chamber (OKOlab). Cell lines were validated by STR testing (Arizona Cancer Center EMSR core facility) and screened for mycoplasma (Biotool).

#### 2D HYDROGELS AND 3D CONDITIONING

For mechanical preconditioning, cells were cultured on 2D polyacrylamide, collagen I-conjugated hydrogels, lab-made or purchased (Petrisoft, Matrigen). Soft hydrogels were either 0.5 kPa (Petrisoft) or <1.0 kPa (lab-made), while stiff hydrogels were either 8.0 kPa (Petrisoft) or lab-made (7.0-8.0 kPa). Prior to making hydrogels, glass coverslips were first pre-treated with a 2% solution of 3-aminopropyltrimethoxy silane (Sigma) in isopropanol, for 10 minutes. After washing, coverslips were treated with 1% Glutaraldehyde for 30 minutes, washed, and dried. To generate the hydrogels, a final ratio of 3/0.055% and 5/0.5% acrylamide/bis-acrylamide were used for <1.0 kPa and 7-8 kPa hydrogels, respectively. Acrylamide and bis-acrylamide were diluted in 50 mM Hepes (pH 8.5), with 0.1% APS and 0.2% TEMED added to the gel solution. Following polymerization on pre-treated glass coverslips, hydrogels were stored at 4°C in PBS until prepared for matrix coating. For matrix coating, hydrogels were treated with 2 mg/ml Sulfo-SANPAH (Life Technologies) and were placed under long wavelength UV light for 5 minutes. Hydrogels were then washed with PBS and coated with 30 μg/ml rat-tail collagen I (Corning) for 1 hour at 37°C, and then washed with PBS again prior to plating. Cells were fed every 2 days and split/assayed at ~80% confluence. Long-term viability was confirmed with LIVE/DEAD Viability/Cytotoxicity (Thermo Fisher) *in situ* and DAPI exclusion during flow cytometry. For 3D conditioning, cells were grown for 7 days in 1.0 mg/mL rat-tail collagen-I (soft), or 1.0 mg/mL rat-tail collagen-I crosslinked with 0.0175% PEG-di(NHS) (MP Biomedicals) (stiff). Cells were spun down and collected after 20 min incubation in Collagenase type I (0.25%) (Stemcell Technologies). Hydrogel stiffness was verified by AFM (W.M. Keck Center for Surface and Interface Imaging).

#### VECTORS AND VIRUS PRODUCTION

The plasmid pCMV-msRUNX2 (a gift from Gerard Karsenty, Columbia) was used as the source for RUNX2 cDNA, and ERK-target site mutants were made using Quikchange (Agilent): RUNX2-

S301A-S319A (**RUNX2-SA**) which is unresponsive to ERK stimulation, and RUNX2-S301E-S319E (**RUNX2-SE**) which exhibits high basal transcriptional activity in the absence of ERK stimulation[1]. RUNX2-WT, RUNX2-SA and RUNX2-SE were then subcloned into lentiviral transfer plasmid pCIG3 (Addgene #78264, a gift from Felicia Goodrum, which was first modified to express a puromycin resistance gene in place of GFP). These mutants were used for *in vitro* and *in vivo* analyses. For human RUNX2 overexpression we subcloned RUNX2-I (MRIPV isoform, GeneCopoeia #EX-I2457-Lv105) into pCIB (Addgene #119863), and the human-equivalent ERK-target sites were made using Quikchange and subcloning: wild-type pCIB-hsRUNX2, pCIB-hsRUNX2-S280A-S298A and hsRUNX2-S280E-S298E. These were used for additional *in vitro* validations. For live-cell actin dynamics, we used pLenti Lifeact-iRFP670- BlastR (Addgene #84385). For constitutive activation of MAPK pathway, we used pBabe-Puro-MEK-DD (a gift from William Hahn, Addgene #15268). shRNA for RUNX2 were purchased from Dharmacon: shRUNX2#01 V2LHS_15065 (TCTGGAAGGAGACCGGTCT); shRUNX2#02 V2LHS_223856 (TACAAATAAATGGACAGTG). For virus production, HEK293T cells were transfected at 60% confluence using Fugene HD (Promega) in OptiMEM (Corning) with transfer plasmid and second-generation lentiviral packaging system (psPAX2 and pMD2.G, Addgene #12260 and #12259, gifts from Didier Trono) or pCL-Ampho (Novus) for lentiviral or retroviral production, respectively. Virus was collected 48-72 hours post-transfection, clarified by 0.45 µm filters. Recipient cells were infected at 50% confluence with virus at a 1:1 dilution with culturing media and polybrene (10µg/mL). Puromycin selection was started 48 hours post-infection.

#### CYTOSKELETAL DYNAMICS

After mechanical preconditioning, SUM159, SKBR3 and MS-SKBR3.1 cells expressing iRFP-LifeAct were trypsizined from their hydrogels and plated onto No. 1.5 glass MatTek dishes which had been pre-treated overnight with DMEM + 10% FBS, and then incubated for 10 hours to ensure maximal spreading before analysis (verified by size equilibrium). Cytoskeletal dynamics score was obtained by automatic tracing of iRFP signal and averaging single-cell displacement over three sequential 1 hour intervals. Imaging was acquired with a 20X Plan Apo 0.75 N objective (Nikon) and an ORCA-Flash 4.0 V2 cMOS camera (Hamamatsu).

#### MULTIDIMENSIONAL TRACTION FORCE

Cells were mechanically-preconditioned as indicated, and then plated onto 1.7 kPa or 8.5 kPa bead-embedded collagen I-coated hydrogels (30 µg/ml), prepared as detailed above. Imaging commenced 4 hours after plating onto bead-embedded hydrogels that were either 1.7 kPa or 8.5 kPa. Fluorescent microsphere beads (0.5 µm; Life Technologies) were dispersed throughout the hydrogels and excited with a red HeNe diode (561 nm) laser. PKH67-stained cells were visualized with an Argon (488 nm) laser. Three-dimensional image stacks were acquired using a Nikon A-1 confocal system mounted on a Ti-Eclipse inverted optical microscope controlled by NIS-Elements Nikon Software. A Plan Fluor 40X air 0.6 N objective (Nikon) mounted on a piezo objective positioner was used, which allowed imaging speeds of 30 frames per second using a resonant scanner. Confocal image stacks of 512 × 512 × 128 voxels (108 × 108 × 38 $\mu m^3$) were recorded every 30 min with a z-step of 0.30 µm. Cell-induced full-field displacements were measured as previously described[2] using the FIDVC algorithm[3].

#### INVASION ASSAY

The invasion assays were modified from Padilla-Rodriguez et al[4]. Briefly, for SUM159 experiments, 75,000 preconditioned single cells were suspended in a dome of 15 µL Matrigel (Corning), spotted onto silanized 8-well coverslip chamber slides (LabTek), incubated for 30 min, and then embedded in 1 mg/mL neutralized rat tail collagen-I (Fisher) crosslinked with 0.0125% PEG-di(NHS) (MP Biomedicals) (see Fig. 1a). Imaging was performed in a 16 hours period, starting 18 hours after embedding. For SKBR3 and MS-SKBR3.1 experiments 180,000 cells were suspended and embedded as above, and then imaging was performed immediately for 24 hours. Invaded cells which divided during the imaging periods were counted as one cell in order to mitigate any differences in proliferation amongst experimental groups. Invasion front translocation was calculated by tracking the midpoint of the cluster of cells at the Matrigel:collagen interface which align perpendicular to the direction of movement using DIC cell tracking in Elements software (Nikon). Imaging was acquired with a 20X Plan Apo 0.75 N objective (Nikon) and an ORCA-Flash 4.0 V2 cMOS camera (Hamamatsu).

#### RNA-SEQ LIBRARY PREPARATION, SEQUENCING, AND NORMALIZATION

SUM159 cells were cultured for 2 weeks on collagen I-conjugated hydrogels that reflect native human breast tumor stiffness corresponding to regions of high cellularity/low matrix deposition (0.5 kPa; Petrisoft, Matrigen), versus low cellularity/high matrix deposition (8.0 kPa; Petrisoft, Matrigen)[5]. Cells were fed every 2 days and were split or analyzed when ~80% confluent. 3 biological replicates of each stiffness were processed for RNA extraction using Isolate II RNA kit (Bioline), and 1µg from each sample was used for polyA selection with [Oligo d(T) Magnetic Beads,New England BioLabs #S1419S]. mRNA was converted into sequencing libraries as previously detailed[6]. In brief, RNAs were fragmented and ligated to barcoded adapters (Bioo Scientific, NEXTflex DNA Barcodes). Quantitative RT-PCR was used to determine the optimal number of cycles to amplify each library as to achieve sufficient DNA quantity while maintaining the diversity of the library (10-20 cycles). Libraries were then amplified, and fragments with insert sizes between 225 and 375 bp were isolated by gel purification, pooled at equimolar concentrations, and submitted for massively parallel high-throughput sequencing (single end, 50 base pairs) on a Hi-Seq4000 (Illumina) at the University of Chicago's Genomics Core using manufacturer protocols.

#### RNA-SEQ DIFFERENTIAL EXPRESSION, PATHWAY ENRICHMENT, AND UPSTREAM REGULATOR ANALYSES

De-multiplexed fastq files were mapped to the human transcriptome (hg38) in STAR[7] using default settings and organized into tag directories using makeTagDirectory in the HOMER[8] software suite. The number of mapped, uniquely aligned reads suggested good coverage of the transcriptome and diversity in the sample set. Hierarchical clustering recapitulated that samples clustered by group membership, as expected, confirming that differences in transcriptomes were driven by cellular matrix environments.

Differential gene expression was calculated in DESeq[9] using a 5% False Discovery Rate (FDR) as defining the differential gene set. Pathway enrichment analysis was performed in Metascape[10] using additional 4-fold cutoffs to define up- and down-regulated genes between soft and stiff samples. Upstream Regulatory analysis was performed on differentially expressed genes using a more inclusive 2-fold cutoff with Ingenuity Pathway Analysis (IPA) software (Qiagen), which returns a list of genes having enriched curated connections to the input gene set as a method to predict upstream regulators. Candidate regulators were restricted to genes having receptor or transcription factor function because these provide a straightforward mechanism for how the mechanical stiffness signal may become integrated in cells to cause differential gene expression. Candidate upstream regulators with prediction $P < 0.05$ were exported and intersected with the gene set annotated as “Metastasis Associated Genes” from the Human Cancer Metastasis Database[11]. From this list of intersecting genes, we then manually searched the literature for those with gene bookmarking function or “transcriptional memory” association. We do not exclude the possibility that we may have missed some epigenetic memory-associated genes that are not well curated. For gene ontologies associated with the stiffness-induced gene set, we also used Enrichr[12], through query of the Human Phenotype Ontology[13] and MGI Mammalian Phenotype[14] libraries.

#### MECHANICAL CONDITIONING (MeCo) SCORING AND PATIENT DATA ANALYSIS

The initial gene set for MeCo scoring was derived from RNA-seq differential expression between SUM159 cells grown on stiff vs soft hydrogels for 2 weeks. Genes that had $P_{adj} < 0.05$ and $|\log_2 FC| > 1$ were considered differentially expressed (FC = fold change). After removing genes associated with proliferation[15], there were a total of 3,822 remaining differentially expressed genes. Out of these genes, 1,143 had a positive $\log_2$FC while 2,679 had a negative $\log_2$FC. Genes with a positive $\log_2$FC were considered to be associated with stiffness, while genes with a negative $\log_2$FC were associated with softness.

We queried three microarray studies from the Gene Expression Omnibus (GEO) to test the clinical association between mechanical conditioning and bone metastasis: GSE2034, GSE2603, GSE12276. Studies GSE2034 and GSE2603 were sequenced using the Affymetrix Human Genome U133A array, and study GSE12276 was sequenced using the Affymetrix Human Genome U133 Plus 2.0 array. The normalized expression matrix was downloaded from GEO using the R GEOquery package and all values were $\log_2$ transformed. In addition, breast tumor samples that underwent sequencing in multiple GEO studies were treated as a single sample [16]. After merging the studies GSE2034, GSE2603, GSE12276 together, there were 12,403 overlapping genes. Out of these 12,403 genes, 2,210 genes were in common with the 3,822 RNA-seq differentially expressed genes. 711 out of the 2,210 genes were associated with stiffness and 1,409 out of the 2,210 were associated with softness (Extended Data Fig. 3a). In conjunction, we used the METABRIC 2019 (molecular dataset) in our analysis to act as an independent study. There were 2,949 genes that overlapped with the RNA-seq gene signature and the METABRIC dataset (942 stiff-associated and 2,007 soft-associated) (Extended Data Fig. 3b).

MeCo score calculation for each patient was performed by taking the average gene expression differences between stiff- and soft-associated genes: [mean expression (stiff genes) – mean expression (soft genes)]. Unlike gene signatures normally used to define cancer subtypes, the MeCo score is patient-specific, so the expression profiles of other samples within the same study do not affect the MeCo score. Moreover, because the MeCo signature subtracts normalized contributions from two sets of genes, any chip- or batch-specific effect that is gene-independent will automatically cancel, making it more robust and transferable across studies[17]. Thus, we combined MeCo scores from the GSE2034, GSE2603, GSE12276 studies and were able to increase the power of our analysis. Notably, we are assuming that the Affymetrix Human Genome U133A array and the Affymetrix Human Genome U133 Plus 2.0 array share similar probe affinities for genes that are in common between arrays.

To optimize the utility of the MeCo score, we refined the RNA-seq gene signature by identifying the overlapping genes that are associated with stiffness or softness, and that are positively and negatively associated with bone metastasis in the three GEO studies described above. The R package *limma* was used to calculate $\log_2$FC between bone metastasis positive patients and bone metastasis negative patients, while controlling for study and subtype in a linear regression framework. Tumor subtypes were identified using the PAM50 signature from the R package *genefu*. This analysis produced 1,051 genes upregulated with bone metastases and 1,069 genes downregulated with bone metastases. Of the 1,051 upregulated genes, 323 were associated with stiffness. Of the 1,069 down regulated genes, 681 were associated with softness. In total, there are 1,004 genes in the refined MeCo score. Furthermore, 919 genes from this refined MeCo score overlapped with the genes represented in the METABRIC expression study and were used to reassess overall survival in that cohort.

The genes used for MeCo score calculations are listed in Supplementary Information (Tables 2-4). For proliferation scoring, we used the normalized, average gene expression of the proliferation-associated genes[15] listed in Supplementary Information (Table 5).

Two independent datasets were used to validate the $MeCo^{refined}$ score: the NKI dataset from van de Vijver et al. 2002[18] and the METABRIC 2019[19]. Note that for the bone metastasis-free survival (BMFS) analysis using the METABRIC 2019, out of the patient subset with gene expression data (METABRIC molecular dataset), we selected all patients with complete recurrence history (Complete.Rec.History=YES). In this analysis, patients with no bone metastasis were censored using their TDR data, which is time until last follow-up or distant relapse; patients with no distant relapse (DR=0) were assumed to not having bone metastasis. Time-to-bone-metastasis (TTBM) analysis was performed using all patients with bone metastasis.

Furthermore, we wanted to test whether mechanical conditioning contributes significantly to the power of the $MeCo^{refined}$ score, or whether our results are simply driven by the gene expression patterns observed in bone metastasis positive and negative tumors. Motivated by the methodology in Venet et al. 2011[20], we generated 1,000 matched random gene sets and compared their performance against $MeCo^{refined}$. Each gene set initially consisted of 2,120 randomly selected genes to mimic the original MeCo gene set. To simulate the calculation of the $MeCo^{refined}$ score for each of the 1,000 random gene sets, we used the same linear regression analysis between bone metastasis positive and bone metastasis negative samples as before. For each gene set, the top 1,004 genes were ranked and separated based on positive $\log_2$FC and negative $\log_2$FC from the regression analysis. Positive genes were considered to be associated with stiffness and negative genes were associated with softness. The randomized versions of the refined MeCo scores were calculated by taking the difference between the mean gene expression of stiff genes and the mean gene expression of soft genes. Distributions of the log rank statistic of BMFS and TTBM for the randomized gene sets were created using the combined cohort of 560 patients. The log rank statistics for both BMFS and time-to-BM computed from the true $MeCo^{refined}$ gene signature were significantly higher than expected from matched random gene sets ($P < 0.05$ and $P < 0.001$).

#### THE ASSAY FOR TRANSPOSASE ACCESSIBLE CHROMATIN ('ATAC-SEQ')

We collected chromatin accessibility data genome-wide across a 7-day time course of transitioning cells from stiff to soft substrates using the assay for transposase accessible chromatin ('ATAC-seq'). ATAC-seq was performed on 50,000 SUM159 cells using the previously described protocol[21]. Libraries were run on a 10% TBE gel and DNA from 175-225 bp was extracted for sequencing. For each time point, triplicate experiments were conducted to allow for quantitative comparisons of accessibility across the time course. After generating libraries, samples were equimolar pooled and sequenced on an Illumina NextSeq High output run (single-end 75bp). After filtering out poor samples based on basic quality control, we retained triplicates for 12 hours after transition (St7/So0.5), 1 day (St7/So1), 2 days (St7/So2), and 5 days (St7/So5), and 2 replicates for the 7-day time point (St7/So7); the third replicate from this time point was excluded because of low sequencing depth (<300,000 unique, autosomal mapped reads vs >6,000,000 for all other samples) and low fraction of reads in peaks called on the sample (0.24 vs 0.39-0.63 for all others). In addition, we retained triplicates for two sets of control samples that were maintained on stiff substrate for 1 day and 7 days after the initial 7-day preconditioning on stiff substrate (St7/St1 and St7/St7, respectively). We then identified peaks of accessibility (also called "hypersensitive sites") on each sample to generate a genome-wide map of accessible regulatory elements using MACS2[22]. To compare the similarity of peaks identified in each time point, we used BEDTools[23] to calculate all pairwise Jaccard indices and generated a heatmap with the 'heatmap.2' function in the 'gplots' package in R. To allow for quantitative comparisons in accessibility between time points we generated a master list of peaks, taking the union of all peaks identified in each of the time points and then counted how many reads mapped to each peak for each sample. These values were normalized for read depth by dividing by how many million reads were contained in the union peak set for each sample. Finally, the read depth-normalized values were $\log_{10}$-transformed (after adding a small constant) and median normalized. Principal component analysis of this matrix confirmed that time was a major predictor of quantitative differences in accessibility for sites common to all samples. A likelihood-ratio test framework was used to identify sites that were differentially accessible in one of the soft matrix time points relative to all of the stiff matrix controls:

$$Y_{ij} = \mu_i + \beta_i X_j + \varepsilon_{ij}$$

$Y_{ij}$ represents the accessibility of site i for sample j, $\mu_I$ is the mean accessibility for site i, $X_j$ is the status of sample j ("soft" vs. "control"), $\beta_i$ is the effect of time spent on soft substrate on accessibility of site i, and $\varepsilon_{ij}$ is an error term. For each site, we compared a model with a "soft vs control" term to a nested model with just an intercept using a likelihood-ratio test. P-values were adjusted for multiple comparisons using the q-value calculation in the 'qvalues' package in R[24]. For this analysis we focused particularly on quick changing *versus* delayed changing sites. Sites that we characterized as 'quick' are those that exhibited differential accessibility at 12 hours after transitioning to soft substrate; we identified 7,607 sites that became more accessible and 2,430 sites that became less accessible at 12 hours (at a false discovery rate or 'FDR' of 1%). Sites that we characterized as 'delayed' are those that did not change for the first 2 days but then changed accessibility by days 5 and 7. To determine the delayed sites, we first identified sites that were significantly differentially accessible by day 5 and 7 (at an FDR of 1%) and then excluded any sites that were also identified at any of the earlier time points (at a relaxed FDR of 10%); this analysis yielded 9,816 sites that became more accessible and 7,277 sites that became less accessible. Importantly, the delayed sites would include regulatory elements for genes exhibiting transcriptional memory.

#### PATHWAY ENRICHMENT ANALYSIS OF ATAC-SEQ DATA

In order to ascertain whether differentially accessible sites for each of the dynamic patterns (quick and delayed closing) were enriched near genes in specific pathways we used the Genomic Regions Enrichment of Annotations Tool ('GREAT'[25]). For both the quick closing and delayed closing sites (defined above), we uploaded a bed file of all sites passing the respective thresholds to great.stanford.edu and used the whole genome as the background to look for ontology enrichments using the default settings.

#### MOTIF ENRICHMENT ANALYSIS

*De novo* motif analysis of ATAC-seq-defined regions was performed using the HOMER software suite[26], for subsets of open chromatin regions. Regions unchanged after transitioning to a soft matrix after seven days were open chromatin regions that were not in the 'quick' or 'delayed' closing set. Specifically, enrichments were determined using the findMotifsGenome.pl command with region sizes of 100 basepairs (bp). For quick and delayed changed sites, unchanged sites were used as the background. For unchanged sites, GC-matched 100-bp random genome sequences were used as background.

#### IMMUNOFLUORESCENCE

For RUNX2 localization, cells were preconditioned for 7 days on soft or stiff hydrogels, passaged at ~80% confluence, with media changes every other day. Where drug treatments are indicated, cells were treated with either 1 µg/mL lysophosphatidic acid (LPA; indirect Rho kinase activator; Sigma), 10 ug/mL Rho Activator II (Cytoskeleton Inc.), 20 µM Y27632 (ROCK inhibitor; Sigma), 50 nM jasplakinolide (actin filament stabilizer; Sigma), 1 µM taxol (microtubule stabilizer; Sigma), 30 µM blebbistatin (Myosin II inhibitor; Sigma), 10 µM nocodazole (microtubule destabilizer; Sigma), or 0.1% DMSO for 3 hours in fresh growth media, and then fixed in 4% paraformaldehyde for 20 min at 37°C. Cells were permeabilized in 0.5% TX-100 for 20 min, and blocked for 1H at RT in 5% goat serum + 0.5% BSA in PBS with DAPI (Sigma) and 2% phalloidin-647 (Invitrogen). Cells were then incubated with primary antibodies for 2 hours at RT, and secondary antibodies for 1 hour at RT. The antibodies used were RUNX2 (Sigma Prestige HPA022040 1:250), Tubulin (Sigma T9026 1:250), Human Cytokeratin (Dako clones AE1/AE3 1:250), Paxillin (BD 612405 1:200), phosphoFAK (Thermo 44-625G 1:200), Alexa Fluor goat anti-rabbit 568 1:250 (Invitrogen) and Alexa Fluor goat anti-mouse 488 1:250 (Invitrogen). Samples were mounted in ProLong Diamond Antifade (Thermo-Fisher) and allowed to cure for at least 24 hours before imaging. Image segmentation was performed on the nuclear (DAPI-stained) image for each field. The cytosolic region was defined as a 2.2 µm wide annulus surrounding the nuclear region using Elements software (Nikon).

#### IMMUNOBLOTTING

For mechanotransduction experiments, 250,000 cells were plate on 8.5 cm circular hydrogels, media changed every other day, and passaged at ~80% confluence. Protein lysates were resolved with SDS-PAGE and transferred to nitrocellulose. For ERK immunoblots, the following drugs were added along with fresh media 1 hour before lysis: 20 µM PD98059 (MEK inhibitor; Tocris), 30 µM blebbistatin (Myosin II inhibitor; Sigma), 100 nM dasatinib (Src inhibitor; Tocris) and 1 µM Faki14 (Fak inhibitor; Tocris) or 0.1% DMSO. For mechanical memory extension, cells were preconditioned as above with the following drugs: 2 µM palbociclib (CDK4/6 inhibitor; Sigma), 7 µM decitabine (DNMT inhibitor; Sigma), or 0.1% DMSO. Cells were lysed 36 hours after the last media change in RIPA buffer (1% NP-40, 150 mM NaCl, 0.1% SDS, 50 mM Tris-HCl pH 7.4, 0.5% sodium deoxycholate) supplemented with Halt protease inhibitor cocktail (Pierce) and Halt phosphatase inhibitor cocktail (Pierce). For AKT inhibition cells were conditioned on stiff hydrogels for 7 days with drug changes every other day (1 µM MK-2206; Cayman Chemical). Membranes were blocked in 100% Odyssey Blocking Buffer PBS (LI-COR), incubated with primary antibodies overnight at 4°C in 50% blocking buffer + 50% PBST, and secondary antibodies for 1 hour at RT in 50% blocking buffer + 50% PBST. The antibodies used were RUNX2 (Sigma Prestige HPA022040 1:1000 and Cell Signaling Technology 8486S 1:1000), OPN (Abcam ab8448 1:1000 and Abcam ab91655 1:1000), ERK (Santa Cruz sc-93-G 1:1000), phospho-ERK (Cell Signaling Technology 4377S 1:1000), AKT (Cell Signaling Technology 2920S 1:1000), phospho-AKT (Cell Signaling Technology 4058S 1:1000), Actin (ProteinTech Group 66009-1 1:5,000), Alexa Fluor goat anti-rabbit 680 1:10,000 (Invitrogen) and Alexa Fluor goat anti-mouse 790 1:10,000 (Invitrogen).

#### FLOW CYTOMETRY AND SORTING

Cells were preconditioned on stiff hydrogels for 7 days to encode mechanical memory, and then labelled with CellVue Claret far-red membrane label (Sigma), using 4 µL label in 200 µL Dil C per 1.0 x $10^6$ cells, before transfer to soft hydrogel conditioning for another 7 days. Cells were fed every 2 days and were split/analyzed when ~80% confluent. Cell sorting was performed on a BD FACSAria III using FACSDiva software. The sequential gating strategy is outlined in Extended Data Fig. 7. Compensation was done for each experiment using unstained cells and cells stained with individual fluorophores. After sorting, 100,000 cells from CellVue$^{HIGH}$ or CellVue$^{LOW}$ gates were returned to 22 mm x 22 mm soft hydrogels in order to generate conditioned media for 24 hours, before assaying OPN and GM-CSF protein expression, or 3D invasion. Flow cytometry was performed on a BD FACSCanto II. To quantify OPN expression, 1 hour room temperature incubation with OPN-PE (Abcam ab210835, 1:2500) or IgG control (Abcam ab72465, 1:2500) was used following -20°C 90% methanol fixation/permeabilization and blocking in 10% goat serum for 30 min at RT. Flow cytometry data was analyzed using FlowJo using unstained samples to set gates.

#### QUANTITATIVE REAL-TIME PCR

For RUNX2 knockdown experiments, cells were infected with either GIPZ (non-targeting control), shRUNX2#01 or shRUNX2#02 expressing lentivirus ($n$ = 3 independent virus preparations each) and selected for 1 week with puromycin. 40,000 cells were plated on soft of stiff hydrogels (22 mm x 22 mm) with media changes every other day, and passaged at 80% confluency. For mechanical memory experiments, cells were plated/passaged as above, and cells were lysed 36 hours after last media change. For mechanotransduction drug treatments, cells were treated with either 30 µM blebbistatin (Sigma), 100 nM dasatinib (Tocris), 1 µM Faki14 (Tocris) on collagen I-coated (or 0.1% DMSO on poly-D lysine-coated) hydrogels for 7 days with media/drug change every other day, including the day before analysis. Total RNA was isolated using Isolate II RNA kit (Bioline) and cDNA was then synthesized from 1µg of RNA using XLA script cDNA kit (Quanta BioSciences). Sybr green PCR mix (Bioline) was used for RT-qPCR on the ABI Fast 7500 system Samples were run in triplicates in each experiment and relative mRNA levels were normalized to housekeeping gene EEF1A. Melt curve analysis was performed to verify that each SYBR reaction produced a single PCR product. All SYBR assays were performed using the following PCR cycling conditions: denaturation at 95 °C for 15 min followed by 40 cycles of denaturing at 95 °C for 10 sec, and annealing at 60 °C for 1 min. See Supplementary Information Table 1 for a list of all primers used.

#### COMBINED MECHANORESPONSE ASSAY

All cell lines and patient-derived xenografts were mechanically-conditioned for 36 hours on 0.5 kPa (Petrisoft) or 8.0 kPa (Petrisoft) hydrogels before analysis. Cells were imaged *in situ* using a 10X Plan Apo 0.75 N objective (Nikon) and an ORCA-Flash 4.0 V2 cMOS camera (Hamamatsu). Manual tracing was done to quantify cell spreading on each stiffness. After image acquisition, RT-qPCR analysis of CTGF was performed as detailed above.

#### ENZYME-LINKED IMMUNOSORBENT ASSAY (ELISA)

SUM159 cells were preconditioned for 7 days on stiff hydrogels to encode mechanical memory, and then 500,000 cells were transferred to 8.5 cm soft hydrogels. Media was changed the next day, and then 24 hours after that conditioned media (CM) was collected (2-day-soft CM), spun down to remove cells/debris and snap frozen. Cells were counted and then 500,000 cells were re-plated. This cycle was repeated to generate 4-, 6-, and 8-day soft CM. A GM-CSF Human SimpleStep ELISA kit was used per manufacturer's instructions (Abcam). The colorimetric signal was normalized to total protein lysates from 25% of the cells on the hydrogels after each 2-day conditioning cycle (Coomassie Plus; Pierce). For mechanical memory selection, 100,000 CellVue$^{HIGH}$ or CellVue$^{LOW}$ cells were returned to 22 mm x 22 mm soft hydrogels for 24 hours in order to generate conditioned media (see Flow Cytometry and Sorting), and GM-CSF levels were interpolated from a standard curve per manufacturer's instructions.

#### QUANTIFICATION OF HUMAN CANCER CELLS IN MOUSE TISSUES

Fresh brain, liver and lung tissue was snap frozen and stored at -20°C. Whole organs were pulverized after liquid nitrogen treatment, and 20 mg from each was processed for gDNA using GeneJet genomic DNA purification kit (Thermo).The following previously validated primers were purchased from IDT: Human Alu, Fw: YB8-ALU-S68 5′-GTCAGGAGATCGAGACCATCCT-3′, Rev: YB8-ALU-AS244 5′-AGTGGCGCAATCTCGGC-3′, Probe: YB8-ALU-167 5′-6-FAM-AGCTACTCGGGAGGCTGAGGCAGGA-ZEN-IBFQ-3′[27]. Mouse Actb PrimeTime Std (Mm.PT.39a.22214843.g) was used as an endogenous control to normalize each sample. All TaqMan assays were performed using the same PCR cycling conditions as listed above.

#### SYNTHETIC BONE MATRIX ADHESION AND SPREADING

Osteo Assay surface (synthetic bone matrix) 24-well microplates (Corning) were used. For adhesion, 24 hour-old preconditioned media from corresponding experimental groups were collected, and then 200 µL was pre-absorbed to the Osteo plates for 1 hour prior to adding 1.0 x $10^6$ cells in 100 µL fresh media, and incubated for 30 min. Plates were gently tapped to remove loosely bound cells, and the remaining cells were stained with DAPI and enumerated by microscopy with large-stitch imaging using a 10X Plan Apo 0.75 N objective (Nikon) and an ORCA-Flash 4.0 V2 cMOS camera (Hamamatsu). For spreading measurements on synthetic bone matrix, the above protocol was used except imaging commenced immediately upon addition of cells to the plate. Cell area was quantified at 6 min intervals by manual tracing in Elements software (Nikon), using a 20X Plan Apo 0.75 NA objective (Nikon) and a CoolSNAP MYO CCD camera (Photometrics).

#### OSTEOCLASTOGENESIS *IN VITRO*

Osteoclast precursor RAW 264.7 cells were induced for 4 days in 24-well plates with DMEM + 10% FBS + 50 ng/mL RANKL (Sigma) (growth media), and then cultured with 50% cancer cell preconditioned media (CM) + 50% growth media for another 3 days. CM was collected 24 hours after addition to plates with equal cell counts from each experimental group. Tartrate-resistant acid phosphatase staining was done using the Acid Phosphatase, Leukocyte (TRAP) kit (Sigma) according to manufacturer's instructions. Multinuclear cells that stained positive were enumerated in large-stitched images taken with a 10X Plan Apo 0.75 N objective (Nikon) and a DS-Fi2 color CCD camera (Nikon).

#### *IN VIVO* MODELS

Adult female 6-8 weeks old NOD.Cg-*Prkdc*$^{scid}$ *Il2rg*$^{tm1Wjl}$/SzJ mice (Jax) were randomly allocated into experimental groups. Mice were maintained in pathogen-free conditions and provided with sterilized food and water *ad libitum*. In the intracardiac injection model, 2 x $10^5$ SUM159, SUM159-RUNX2-WT, SUM159-RUNX2-SA or SUM159-RUNX2-SE cells (preconditioned for 7 days on soft hydrogels, stiff hydrogels, or TC plastic, as indicated) were injected (resuspended in 100 µL PBS) into the left cardiac ventricle. Upon harvest, non-osseous tissues were flash frozen for subsequent DNA extraction and human *Alu* quantification. In the intraosseous injection model, SUM159 cells were first preconditioned for 7 days on either soft or stiff hydrogels, then re-plated onto new soft hydrogels for 24 hours prior to 5 x $10^4$ cells (in 5 µL PBS) being injected into the intramedullary space of the right distal femur of each animal. All animal studies were approved by the Institutional Animal Care and Use Committee at the University of Arizona.

#### PATIENT-DERIVED XENOGRAFTS

PDX models were established and gifted by Alana Welm (Huntsman Cancer Institute). For propagation of HCI-005, HCI-011 and HCI-003 tumors, 2 mm x 2 mm frozen chunks were implanted subcutaneously and allowed to grow until their diameter exceeded 1.75 cm and necessitated excision, or the animals showed signs of undue pain or distress. *Ex vivo* analysis of PDX cells was done between p4 and p6. To isolate transformed cells, freshly excised tumor chunks were digested with collagenase/hyaluronidase in DMEM (Stemcell Technologies) for 3 hours at 37°C, using differential adhesion to reduce the proportion of mouse fibroblasts transferred to hydrogels for preconditioning.

#### BIOLUMINESCENCE *IN VIVO*

Once a week, mice were injected with 120 mg/kg luciferin and metastatic dissemination was monitored using AMI X (Spectral Instruments). Mice were killed by $CO_2$ asphyxiation 3-4 weeks after tumor cell injection. Metastatic burden was quantified using AMIView (Spectral Instruments). Exclusion criteria for data analysis were pre-established such that those mice terminated before defined experimental endpoints, for ethical reasons or premature death, were not included in analysis. Investigators were blinded to experimental groups during acquisition of bioluminescence data.

#### X-RAY IMAGING

Mice were anesthetized with 80 mg/kg ketamine to 12 mg/kg xylazine (in a 10 mL/kg volume) and radiographs were obtained (Faxitron). Data were analyzed with ImageJ using pixels$^2$ as the unit of measurement. Investigators were blinded to experimental groups during acquisition and analysis of X-ray data.

#### MICRO-CT IMAGING

Legs were removed from euthanized mice and fixed in neutral-buffered formalin for 24 hours, then dissected free of tissue and scanned on a Siemens Inveon micro-CT at 80 kV with a 0.5 mm filter, using an effective pixel size of 28 microns. The scanned images were reconstructed with Inveon Research Workplace (Siemens) using the Feldkamp algorithm and Shepp-Logan filter. In the intracardiac injection model, bone parameters were determined in the distal femur, starting 3 mm from the growth plate to the top of the epiphysis. In the intraosseous model, cortical bone thickness, volume, and surface area were determined in a 4 mm length of midshaft femur; trabecular bone analysis was excluded in the intraosseous model due to potential destruction caused by the syringe. In order to delineate bone marrow, trabecular bone and cortical bone, signal threshold intervals were set identically for all specimens. All histomorphometric parameters were based on the report of the American Society for Bone and Mineral Research nomenclature[28]. Investigators were blinded to experimental groups during acquisition and analysis of micro-CT data.

#### STATISTICS

Sample sizes were determined based on our previous experience with similar experiments (a minimum of 3 to 5 mice for animal studies, or 2 to 4 biological replicates for *in vitro*/*ex vivo* assays). Statistical significance was assessed with GraphPad Prism 8, using the appropriate tests as listed in each figure legend. For analysis patient survival data, we used Kaplan-Meier plots and the Wilcoxon test.

#### DATA AVAILABILITY

RNA-seq and ATAC-seq data are available at the NCBI Gene Expression Omnibus under accession number GSE127887. All other data are available upon reasonable request.

#### ACKNOWLEDGEMENTS

The authors wish to acknowledge the following: the Experimental Mouse Resource Service, particularly Gillian Paine-Murrieta and Bethany Skovan for their technical assistance; Brenda Baggett for micro-CT imaging; John Fitch, Mark Curry and John Davies for FACS assistance; Michael Whalen and Lindsey Stolze for data processing. This research was supported by NCI grant R01 CA196885-01 (G.M.), NHLBI grant R00 HL123485 (C.R.), NIAID grant R01 AI116629-01A1 (C.F.), the Timothy and Diane Bowden Fellowship (A.W.) and the NCI University of Arizona Cancer Center Support Grant P30 CA023074.

#### CONTRIBUTIONS

A.W. and G.M. designed the study and wrote the manuscript with input from all authors. A.W., M.H., S.P., M.R., B.F., C.G., and R.CP. performed and analyzed cell culture and animal experiments. M.H. performed the traction force experiments while C.F. supervised and contributed intellectually. B.F. performed the X-ray analysis. A.W. and G.M. conceived of the MeCo scoring metric, and A.G., M.P., and G.M. performed iterative refinement. C.R. performed the RNA-seq and upstream regulator analysis. D.C. performed the ATAC-seq analysis. A.G. and G.M. performed the analysis of patient datasets. G.M. supervised the overall study.

#### CONFLICT OF INTEREST

The authors declare no conflict of interest.

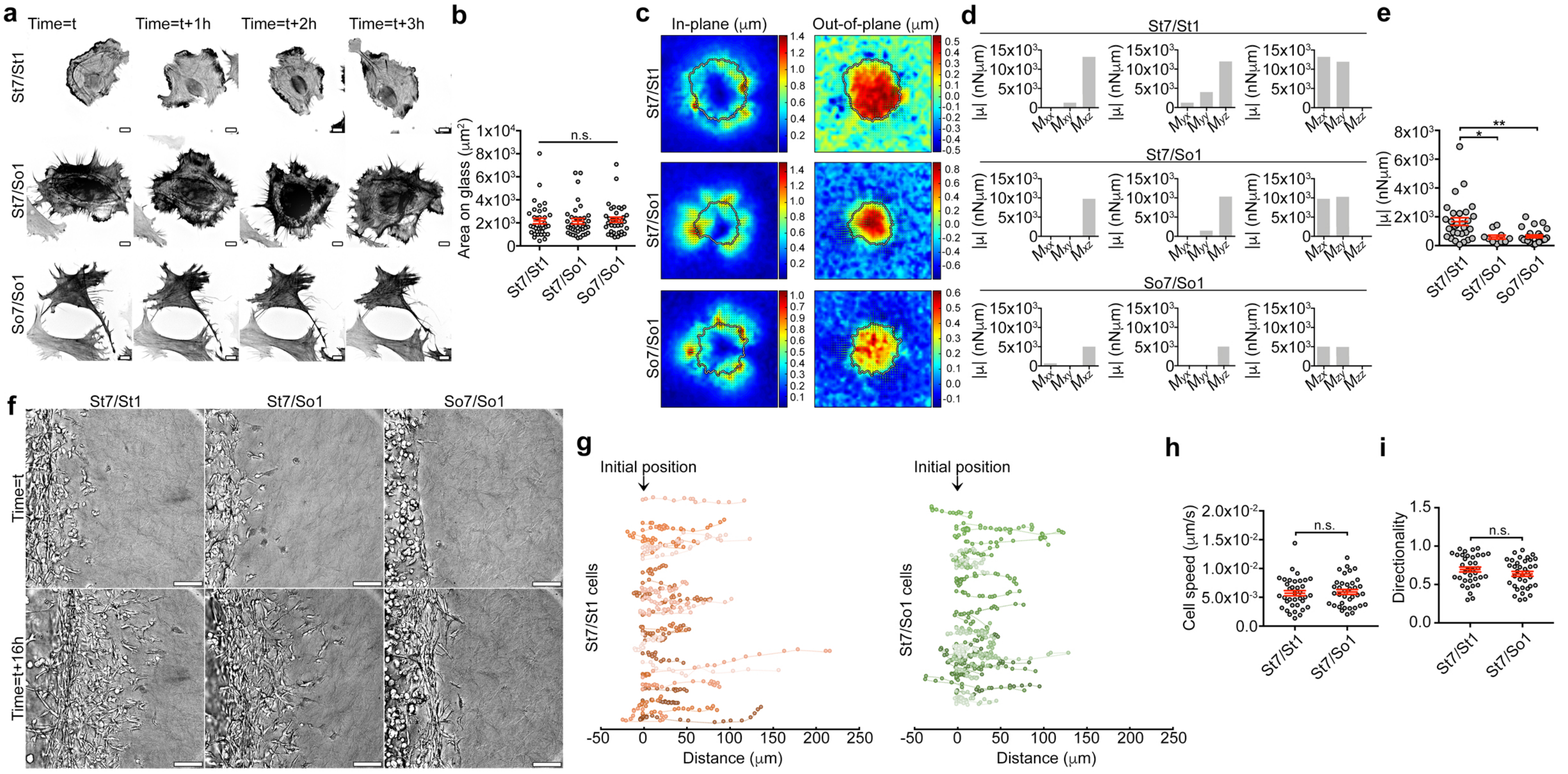


### EXTENDED DATA FIGURE 1

**Extended Data Fig.1. Retention of mechanical memory is not uniform across different cellular dynamics. a,** Cytoskeletal dynamics of iRFP-Lifeact-expressing SUM159 cells related to Fig.1c, scale bars = 10 µm. **b,** Quantification of cell area for cells represented in (**a**) (*n* = 36 cells in each condition from *n* = 3 biological replicates). Data are mean ± s.e.m. *P* is not significant by one-way ANOVA. **c,** Representative heatmaps of 2D traction stress on bead-embedded 8.5 kPa hydrogels of SUM159 cells, preconditioned on stiff and/or soft hydrogels as indicated. **d,** Dipole moment components from representative cells in (**c**). **e,** Quantification of contractile strength on bead-embedded 1.7 kPa hydrogels of SUM159 cells, preconditioned on stiff and/or soft hydrogels as indicated (*n* = 10-29 cells in each condition from *n* = 3 biological replicates). Data are mean ± s.e.m. $^{*}P < 0.05$; $^{**}P < 0.01$, one-way ANOVA with Tukey's multiple comparisons test. **f,** Enhanced depth-of-focus DIC images corresponding to Fig. 1g, scale bars = 100 µm. **g,** SUM159 cell invasion tracks from a representative experiment corresponding to Fig.1g, with 25 min intervals between points. **h,i,** Quantification of cell speed (**h**) and directionality (**i**) from the experiment outlined in Fig. 1g. Data are mean ± s.e.m. *P* is not significant by two-tailed unpaired Student's *t*-test. See Source Data for exact *P* values.

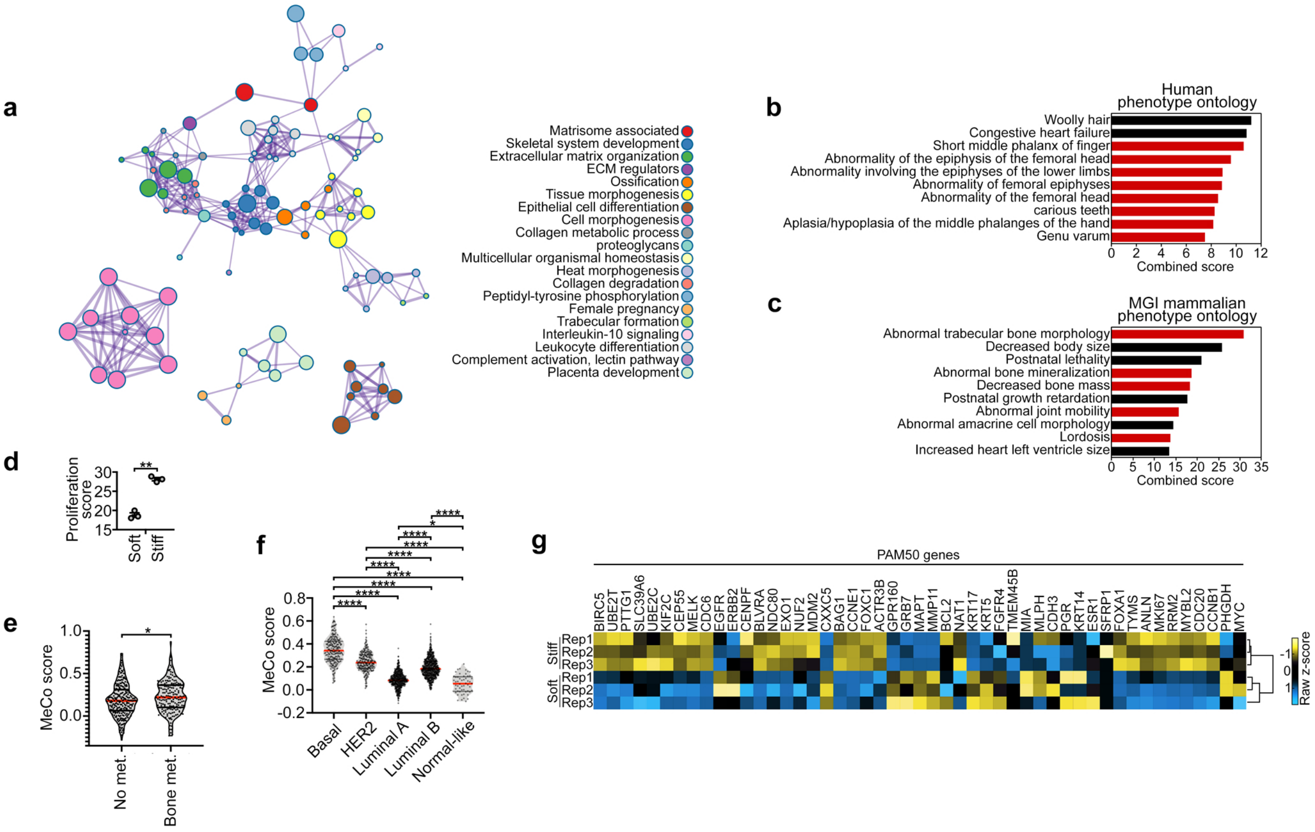


### EXTENDED DATA FIGURE 2

**Extended Data Fig.2. Stiffness-induced genes and mechanical conditioning are associated with skeletal pathologies and bone metastasis. a,** Metascape enrichment network of the stiffness-induced gene set (>4-fold) from SUM159 cells after 2 weeks of mechanical preconditioning. **b,c** Top ten candidate factors from Enrichr analysis of the gene set in (**a**) using the Human Phenotype Ontology library[13] (**b**) and the MGI Mammalian Phenotype library[14] (**c**). Red bars = skeletal pathologies. All hits are significant at $P < 0.05$. **d,** Proliferation score of 2-week soft- and stiff-preconditioned SUM159 cells. Data are mean ± s.e.m. $^{**}P < 0.01$, paired two-tailed t-test. **e,** MeCo scores of patients in the combined cohort in Fig.2d ($n$ = 268 no metastasis, 185 bone metastasis). Data are mean ± s.e.m. $^{*}P < 0.05$, unpaired two-tailed t-test. **f,** MeCo scores for patients in the METABRIC study, stratified by subtype ($n$ = 244 Basal, 243 HER2, 596 Luminal A, 764 Luminal B, 54 Normal-like). Data are mean ± s.e.m. $^{*}P < 0.05$, $^{****}P < 0.0001$, one-way ANOVA with Tukey's multiple comparisons test. **g,** Unsupervised clustering analysis of 2-week soft- and stiff-preconditioned SUM159 cells using the PAM50 gene set. See Source Data for exact $P$ values.

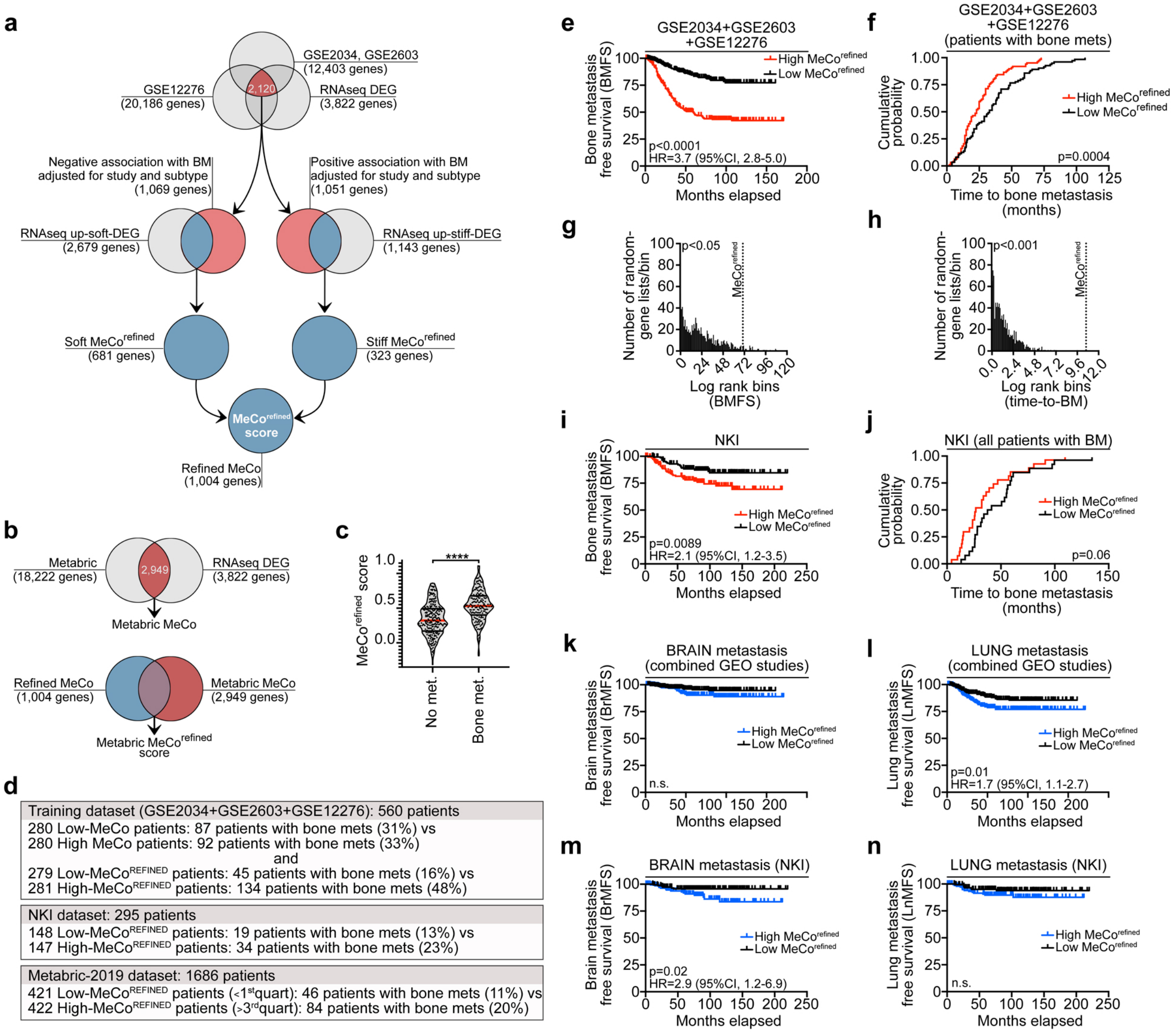


### EXTENDED DATA FIGURE 3

**Extended Data Fig.3. Stiffness-induced genes and mechanical conditioning are associated with skeletal pathologies and bone metastasis. a,** Flowchart of MeCo score refinement, adjusting for study and subtype. **b,** Flowchart of MeCo score refinement for METABRIC analysis. **c,** $MeCo^{refined}$ scores of patients from patients in Fig.2d (*n* = 268 no metastasis, 185 bone metastasis). **d,** Table showing number and percentages of patients with bone metastasis in the analyzed studies. **e,** Kaplan-Meier curve of bone metastasis-free survival in the combined cohort, split at median $MeCo^{refined}$ score (*n* = 281 high $MeCo^{refined}$, 279 low $MeCo^{refined}$). **f,** Time to bone metastasis for patients in (e), split at median $MeCo^{refined}$ score (*n* = 93 high $MeCo^{refined}$, 92 low $MeCo^{refined}$). **g,** Distribution of log-rank statistic of BMFS generated from 1000 random gene sets in comparison with $MeCo^{refined}$ score. **h,** Distribution of log-rank statistic of time-to-BM generated from 1000 random gene sets in comparison with $MeCo^{refined}$ score. **i,** Kaplan-Meier curve of bone metastasis-free survival in the NKI cohort, split at median $MeCo^{refined}$ score (*n* = 147 high $MeCo^{refined}$, 148 low $MeCo^{refined}$). **j,** Time to bone metastasis for patients in (i), split at median $MeCo^{refined}$ score (*n* = 27 high $MeCo^{refined}$, 26 low $MeCo^{refined}$). The median time to metastasis for patients with high MeCo scores was 27 months, compared to 39 months for those with low MeCo scores. **k,** Kaplan-Meier curve of brain metastasis-free survival in the combined cohort, split at median $MeCo^{refined}$ score (*n* = 280 high $MeCo^{refined}$, 280 low $MeCo^{refined}$). **l,** Kaplan-Meier curve of lung metastasis-free survival in the combined cohort, split at median $MeCo^{refined}$ score (*n* = 280 high $MeCo^{refined}$, 280 low $MeCo^{refined}$). **m,** Kaplan-Meier curve of brain metastasis-free survival in the NKI cohort, split at median $MeCo^{refined}$ score (*n* = 147 high $MeCo^{refined}$, 148 low $MeCo^{refined}$). **n,** Kaplan-Meier curve of lung metastasis-free survival in the NKI cohort, split at median $MeCo^{refined}$ score (*n* = 147 high $MeCo^{refined}$, 148 low $MeCo^{refined}$). Kaplan-Meier *P* values calculated with Wilcoxon test. $^{*}P < 0.05$, $^{**}P < 0.01$, $^{****}P < 0.0001$, two-tailed unpaired Student's *t*-test. See Source Data for exact *P* values.

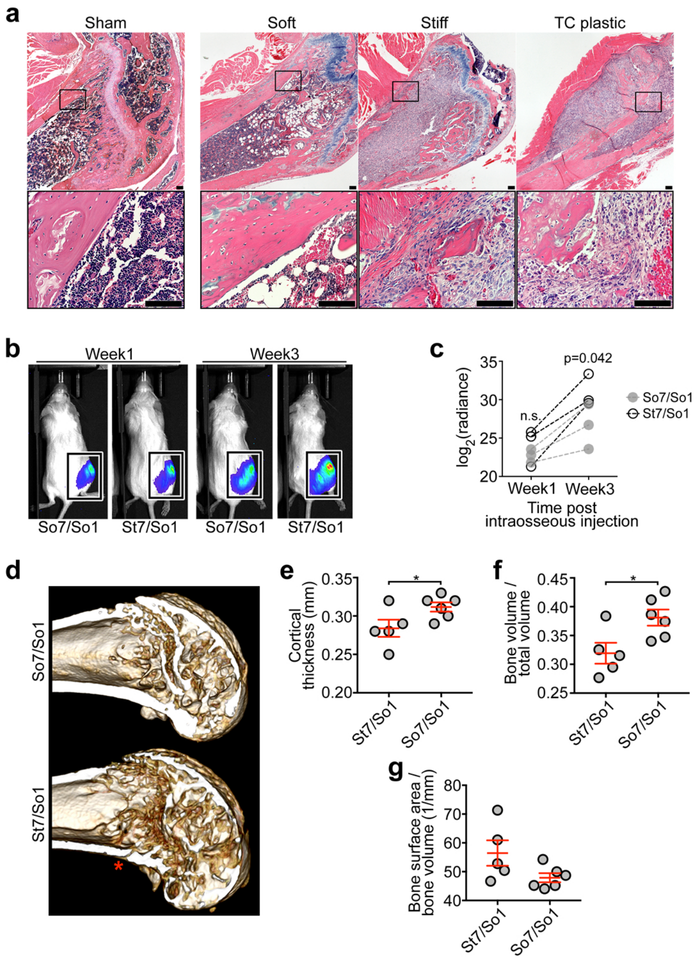


### EXTENDED DATA FIGURE 4

**Extended Data Fig.4. Primary tumor mechanical conditioning and mechanical memory are associated with bone metastasis. a,** H&E staining of femurs from mice bearing 7-day soft-, stiff-, and plastic-preconditioned SUM159 cells, 4 weeks after intracardiac injection. Scale bars = 100 µm. **b,** Bioluminescence images of stiffness-memory (St7/So1) and soft-preconditioned (So7/So1) SUM159 cells after intrafemoral injection, imaged at 1 week and 3 weeks post-injection. **c,** Quantification of bioluminescence in (**b**) ($n$ = 3 mice per group). Two-tailed unpaired Student's $t$-test. **d,** Micro-CT 3D reconstructions of femurs from mice bearing stiffness-memory (St7/So1) and soft-preconditioned (So7/So1) SUM159 cells, 4 weeks after intrafemoral injection. Asterisk marks a fracture. **e-g,** Quantification of cortical thickness (**e**) bone volume/total volume (**f**) and bone surface area/bone volume (**g**) from mice in (**d**) ($n$: mice; St7/So1 5; So7/So1 6). Data are mean ± s.e.m. *$P < 0.05$, two-tailed unpaired Student's $t$-test. See Source Data for exact $P$ values.

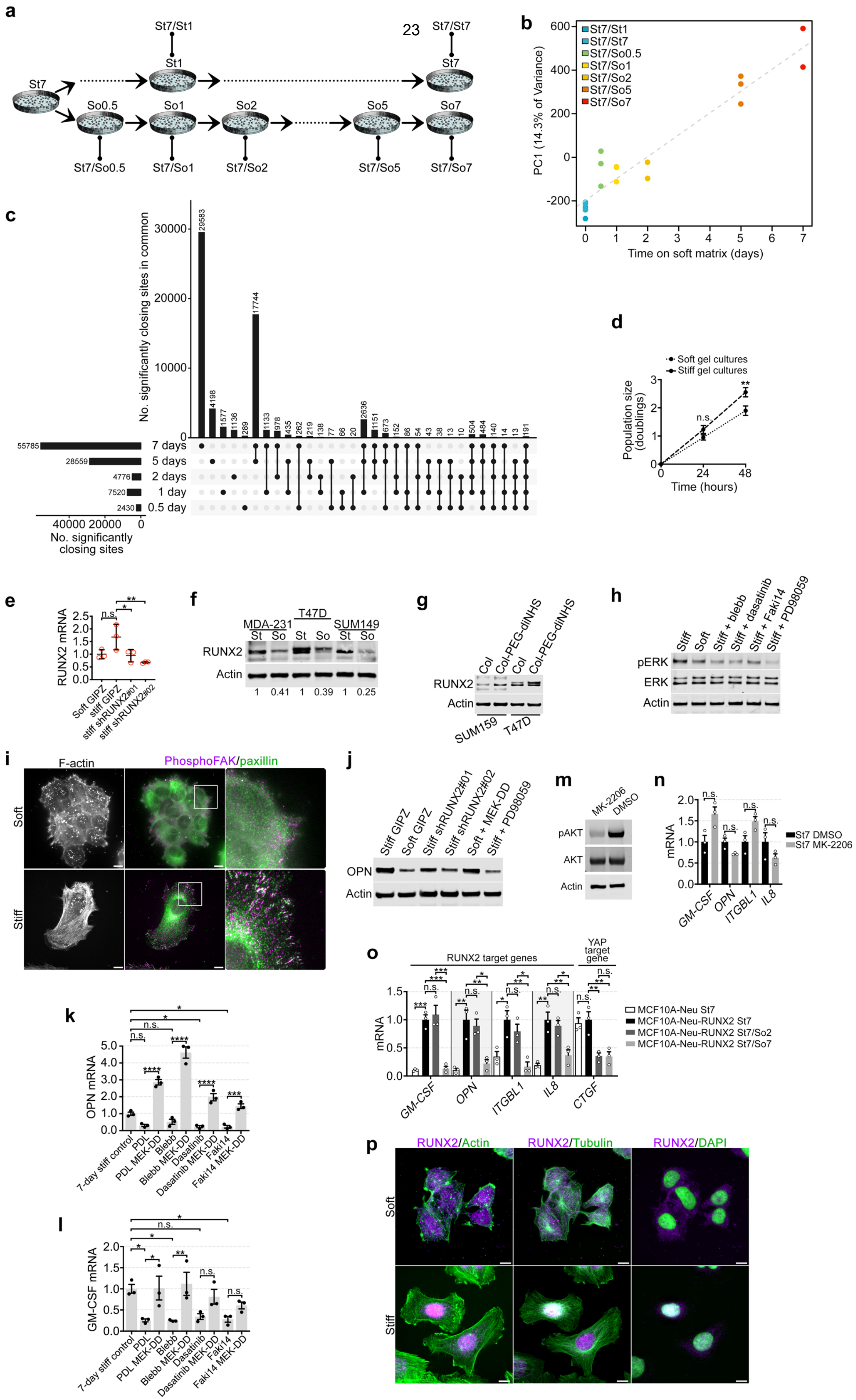


### EXTENDED DATA FIGURE 5

**Extended Data Fig.5. RUNX2 is activated by mechanotransduction. a,** Schematics showing experimental strategy for ATAC-seq experiments; samples were collected at indicated time points after transitioning to either stiff (top row) or soft (bottom row) environments; samples were generated in triplicates. **b,** Principal component analysis (PCA) of ATAC-seq. **c,** Upset plot of the intersection of the different time points. **d,** Population doubling of SUM159 cells at 24 and 48 hours. **P* < 0.05; ***P* < 0.01;****P* < 0.001; *****P* < 0.0001, two-way ANOVA with Sidak's multiple comparisons test. **e,** RT-qPCR of RUNX2 in SUM159 cells preconditioned for 7 days on soft and stiff hydrogels with non-targeting shRNA (GIPZ), or on stiff hydrogels with two shRNAs targeting RUNX2 (*n* = 3 biological replicates). **P* < 0.05; ***P* < 0.01;****P* < 0.001; *****P* < 0.0001, one-way ANOVA with Sidak's multiple comparisons test. **f,** Immunoblot of RUNX2 in MDA-231, T47D and SUM149 cells preconditioned on soft and stiff hydrogels for 7 days (representative of *n* = 2 biological replicates). **g,** Immunoblot of RUNX2 in SUM159 and T47D cells cultured for 7 days in 3D matrix consisting of soft 1.0 mg/mL rat-tail collagen-I, or stiff 1.0 mg/mL rat-tail collagen-I crosslinked with PEG-di(NHS) to stiffen the collagen lattice without changing ligand density. (representative of *n* = 3 biological replicates). **h,** Immunoblot of pERK and ERK in SUM159 cells cultured on stiff hydrogels with 20 μM PD98059, 30 μM blebbistatin, 100 nM dasatinib, 1 μM Faki14 or DMSO for 1 hour prior to lysis. (representative of *n* = 3 biological replicates). **i,** Immunofluorescence of pFAK, paxillin, and F-actin in SUM159 cells cultured on stiff or soft hydrogels for 7 days. **j,** Immunoblot of OPN in SUM159 cells preconditioned for 7 days on soft and stiff hydrogels with non-targeting shRNA (GIPZ), on stiff hydrogels with two shRNAs targeting RUNX2, on soft hydrogels with constitutively-active MEK-DD expression, or on stiff hydrogels with MEK inhibitor PD98059 (20 μM). (representative of *n* = 3 biological replicates). **k,l,** RT-qPCR of OPN (**k**) and GM-CSF (**l**) in SUM159 cells preconditioned for 7 days on stiff hydrogels conjugated with either poly D-lysine (PDL) to reduce integrin binding, or collagen-conjugated with DMSO (control), 30 μM blebbistatin, 100 nM dasatinib or 1 μM Faki14 in media changed every other day. (*n* = 3 biological replicates). Data are mean ± s.e.m and normalized to 7-day stiff controls. **P* < 0.05; ***P* < 0.01;****P* < 0.001; *****P* < 0.0001, one-way ANOVA with Sidak's multiple comparisons test. **m,** Immunoblot showing phospho-AKT levels in SUM159 cells treated with DMSO (control) or AKT inhibitor (1 μM MK-2206). (representative of *n* = 2 biological replicates). **n,** RT-qPCR of RUNX2 target genes in SUM159 cells preconditioned for 7 days on stiff hydrogels and treated with DMSO or 1 μM AKT inhibitor MK-2206 (*n* = 3 biological replicates). Data are mean ± s.e.m. Multiple t-test with Holm-Sidak multiple comparisons; adjusted P values are not significant (n.s.). **o,** RT-qPCR of RUNX2 target genes and CTGF (YAP target) in MCF10A-Neu and MCF10A-Neu-RUNX2 cells preconditioned as indicated (*n* = 3 biological replicates). **P* < 0.05; ***P* < 0.01;****P* < 0.001; *****P* < 0.0001, one-way ANOVA with Sidak's multiple comparisons test. **p,** Immunofluorescence corresponding to Fig. 3e. Scale bars = 10 μm. See Source Data for exact *P* values.

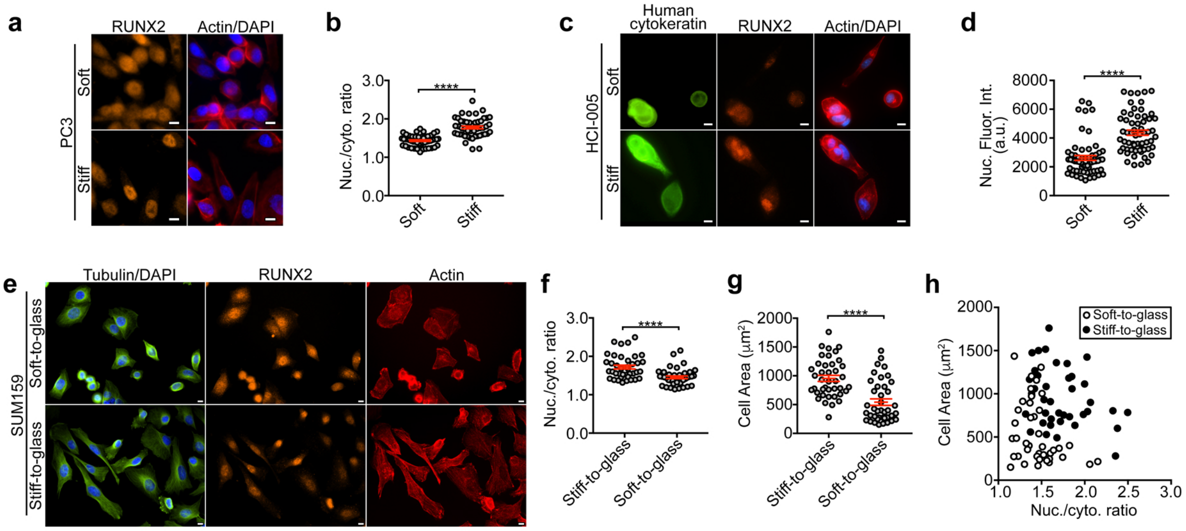


### EXTENDED DATA FIGURE 6

**Extended Data Fig.6. Substrate stiffness *in situ* and stiffness-memory both promote nuclear localization of RUNX2, independent of supraphysiological stiffness tolerance or cell spreading. a,b,** Immunofluorescence staining (**a**) and quantification of nuclear localization (**b**) of RUNX2 in PC3 prostate cancer cells ($n$ = 50 cells each from $n$ = 3 biological replicates). **c,d,** Immunofluorescence staining of RUNX2 (**c**) and quantification of nuclear intensity of RUNX2 in CK+ cells (**d**) in supraphysiological stiffness-naïve patient-derived xenograft HCI-005 tumor cells ($n$ = 60 cells each from $n$ = 3 biological replicates). **e,** Immunofluorescence staining in SUM159 cells, preconditioned on soft and stiff hydrogels for 7 days before transferring to collagen-coated glass for 3 hours. **f,g**, Quantification of nuclear RUNX2 (**f**) and cell area (**g**) from cells in (**e**). **h**, Correlation analysis of (**f,g**) showing no positive intracellular correlation between cell spreading and nuclear RUNX2 in either soft- or stiff-preconditioned cells spreading on glass ($n$ = 40 cells each from $n$ = 3 biological replicates). Soft-to-glass Pearson's r = -0.31 (n.s.); stiff-to-glass Pearson's r = -0.30 (n.s.). Data are mean ± s.e.m. ****$P < 0.0001$, two-tailed unpaired Student's $t$-test. See Source Data for exact $P$ values.

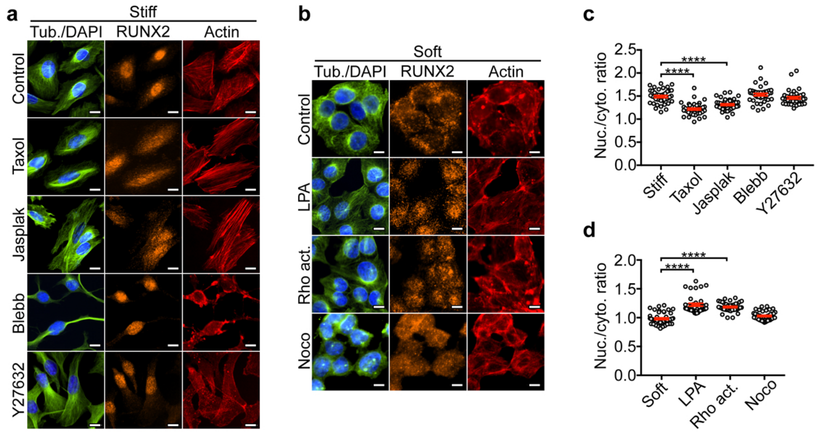


### EXTENDED DATA FIGURE 7

**Extended Data Fig.7. RUNX2 localization is influenced by cytoskeletal dynamics. a,** Immunofluorescence staining of RUNX2 in SUM159 cells preconditioned for 7 days on stiff hydrogels, treated with DMSO (control), 1 μM taxol, 50 nM jasplakinolide, 30 μM blebbistatin or 20 μM Y27632, added 3 hours before fixation. Scale bars = 10 μm. **b,** Immunofluorescence staining of RUNX2 in SUM159 cells preconditioned for 7 days on soft hydrogels, with DMSO (control), 1 μg/mL lysophosphatidic acid (LPA), 10 μg/mL Rho Activator II, or 10 μM nocodazole, added 3 hours before fixation. Scale bars = 10 μm. **c,** Quantification of (**a**) ($n \geq 40$ cells each condition from $n = 3$ biological replicates). **d,** Quantification of (**b**) ($n \geq 40$ cells each condition from $n = 3$ biological replicates). Data are ± s.e.m. ****$P < 0.0001$, one-way ANOVA with Dunnett's multiple comparisons test. See Source Data for exact *P* values.

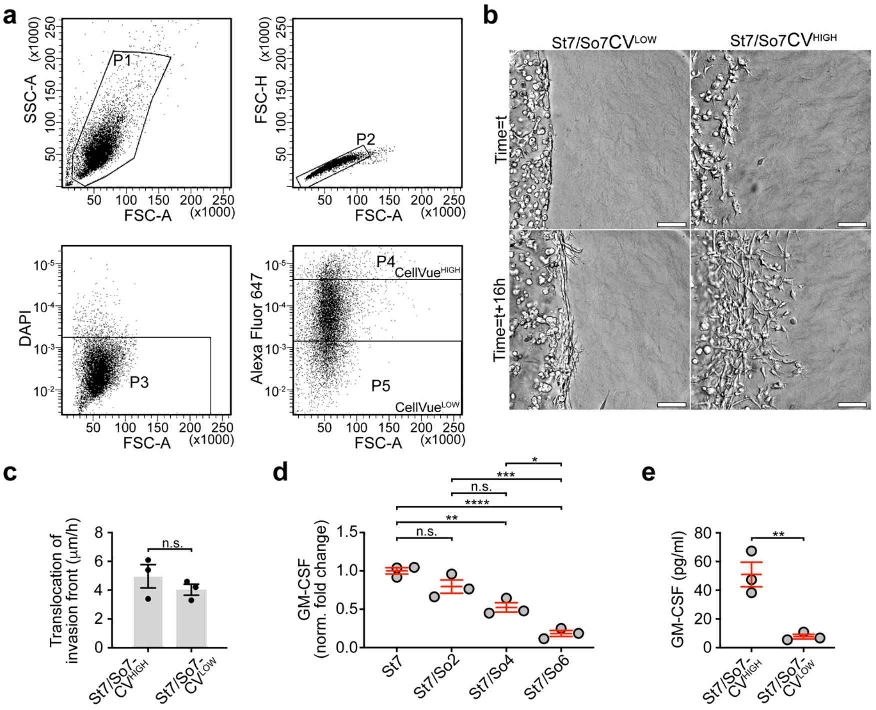


### EXTENDED DATA FIGURE 8

**Extended Data Fig.8. Mechanical memory is more durable in low-proliferative cells. a,** FACS plots depicting the sequential gating strategy for enriching single, live (by exclusion of DAPI), CellVue Claret Far-Red (Alexa Fluor 647) HIGH (P4) and LOW (P5) SUM159 cells. CellVue dye retaining cells, St7/So7CellVue$^{HIGH}$, proliferated less than St7/So7CellVue$^{LOW}$ cells. **b,** Enhanced depth-of-focus DIC images corresponding to Fig.3i. **c**, Rate of translocation of the invasion front corresponding to Fig.3i (*n* = 3 biological replicates with *n* = 3 technical replicates). **d,e,** GM-CSF ELISA for SUM159 cells preconditioned on stiff and/or soft hydrogels as indicated (*n* = 3 biological replicates with *n* = 3 technical replicates). Data are mean ± s.e.m. *$P < 0.05$; **$P < 0.01$;***$P < 0.001$; ****$P < 0.0001$, one-way ANOVA with Tukey's multiple comparisons test for (**d**), **$P < 0.01$, two-tailed unpaired Student's *t*-test for (**c**,**e**). See Source Data for exact *P* values.

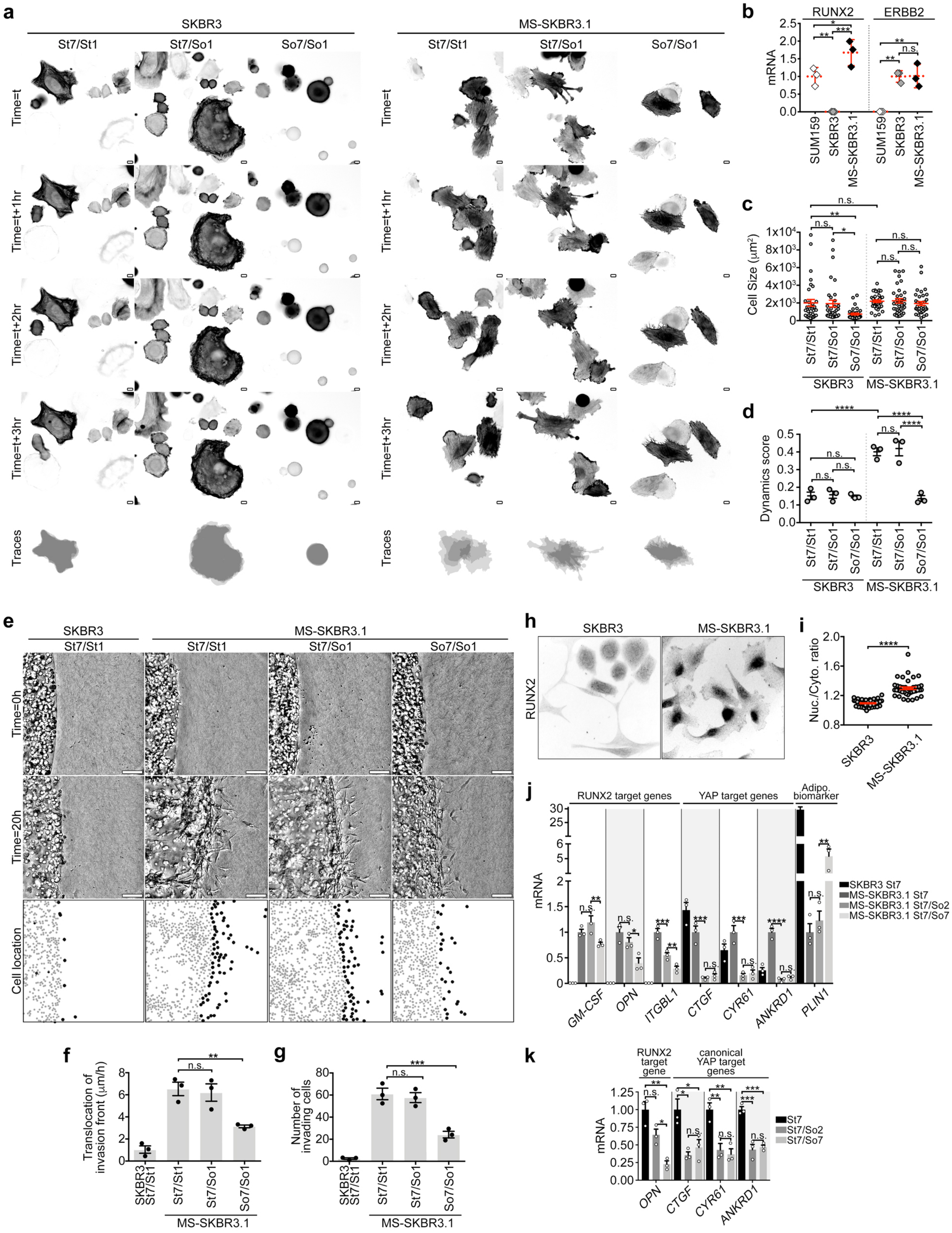


### EXTENDED DATA FIGURE 9

**Extended Data Fig.9. Mechanically-sensitized MS-SKBR3.1 cells have functional mechanical memory and increased nuclear localization of RUNX2. a,** Cytoskeletal dynamics of iRFP-Lifeact-expressing SKBR3 and MS-SKBR3.1 cells, 10 hours after plating on collagen-coated glass, preconditioned on stiff and/or soft hydrogels as indicated. Scale bars = 10 µm. See Supplementary Movie 4. **b,** RT-qPCR of RUNX2 and ERBB2 in cells preconditioned for 7 days on stiff hydrogels (*n* = 3 biological replicates). Data are ± s.e.m. **P* < 0.05; ***P* < 0.01;****P* < 0.001, one-way ANOVA with Tukey's multiple comparisons test. **c,** Quantification of cell size from (**a**), showing differences amongst the SKBR3 cell groups but not MS-SKBR3.1 cell groups (*n* = 36 cells total in each condition from *n* = 3 biological replicates). ). Data are mean ± s.e.m. **P* < 0.05; ***P* < 0.01, one-way ANOVA with Holm-Sidak's multiple comparisons test. **d,** Quantification of dynamics score from (**a**) showing differences amongst the MS-SKBR3.1 cell groups but not SKBR3 cell groups (*n* = 36 cells total in each condition from *n* = 3 biological replicates). Data are mean ± s.e.m. *****P* < 0.0001, one-way ANOVA with Holm-Sidak's multiple comparisons test. **e,** Invasion fronts of SKBR3 and MS-SKBR3.1 cells, preconditioned on stiff and/or soft hydrogels as indicated, after 20 hours of live-cell tracking in 3D collagen. Gray dots = non-invasive cells; black dots = invasive cells. See Supplementary Movie 5. **f,g,** Rate of translocation of the invasion front (**f**) and number of invading cells per field (**g**) from (**e**) (*n* = 3 biological replicates with *n* = 3 technical replicates). Data are mean ± s.e.m. ***P* < 0.01, one-way ANOVA with Tukey's multiple comparisons test. **h,i,** Immunofluorescence staining (**h**) and quantification of nuclear localization (**i**) of RUNX2 in SKBR3 and MS-SKBR3.1 cells (*n* = 36 cells each from *n* = 3 biological replicates). Data are mean ± s.e.m. *****P* < 0.01, two-tailed unpaired Student's *t*-test. See Source Data for exact *P* values. **j,** RT-qPCR of RUNX2 and YAP gene targets and the adipogenic biomarker *PLIN1* in SKBR3 and MS-SKBR3.1 cells preconditioned as indicated (*n* = 3 biological replicates). Data are mean ± s.e.m. **P* < 0.05; ***P* < 0.01; ****P* < 0.001, one-way ANOVA with Tukey's multiple comparisons test. **k,** RT-qPCR of the RUNX2 gene target *OPN*, and three YAP targets, *CTGF*, *CYR61* and *ANKRD1*, in SUM159 cells preconditioned as indicated, and without media-change for 48 hours before sample collection (*n* = 3 biological replicates). Data are mean ± s.e.m. **P* < 0.05; ***P* < 0.01; ****P* < 0.001, one-way ANOVA with Tukey's multiple comparisons test. See Source Data for exact *P* values.

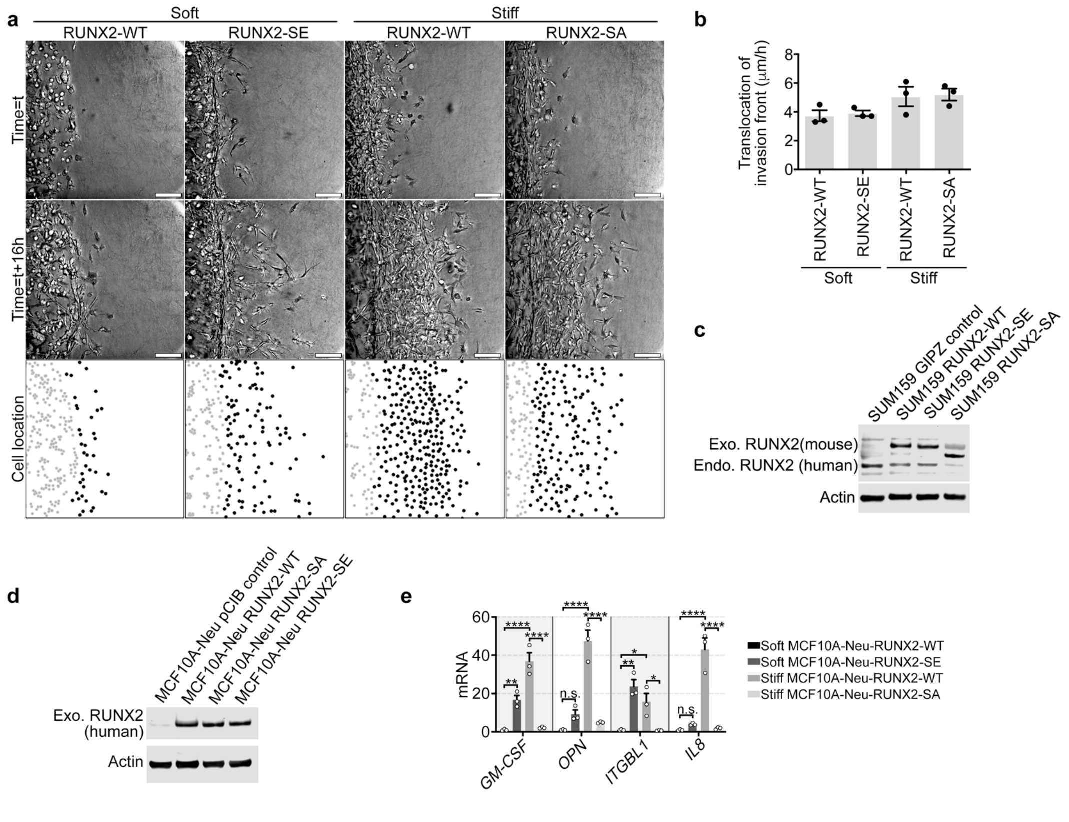


### EXTENDED DATA FIGURE 10

**Extended Data Fig.10. RUNX2-mediated mechanical memory promotes invasion. a,** Invasion fronts of SUM159 cells after 16 hours of live-cell tracking in 3D collagen, Cells overexpressing mouse RUNX2-WT, RUNX2-SA or RUNX2-SE were preconditioned for 7 days on stiff or soft hydrogels, as indicated. See Fig.4a**.** Gray dots = non-invasive cells; black dots = invasive cells. **b**, Rate of translocation of the invasion front from (**a**) (*n* = 3 biological replicates with *n* = 3 technical replicates). Data are mean ± s.e.m. *P* = not significant, one-way ANOVA with Holm-Sidak's multiple comparisons test. **c**, Immunoblot of RUNX2 in SUM159 cells overexpressing GIPZ (control), RUNX2-WT, RUNX2-SE, and RUNX2-SA. (*n* = 3 biological replicates). **d**, Immunoblot of human RUNX2 in MCF10A-Neu cells overexpressing pCIB (control), RUNX2-WT, RUNX2-SE, and RUNX2-SA. (*n* = 3 biological replicates). **e,** RT-qPCR of 4 RUNX2 target genes in MCF10A-Neu cells expressing human RUNX2 WT or mutants preconditioned for 7 days on soft and stiff hydrogels (*n* = 3 biological replicates). Data are mean ± s.e.m. **P* < 0.05; ***P* < 0.01; ****P* < 0.001, one-way ANOVA with Tukey's multiple comparisons test. See Source Data for exact *P* value.

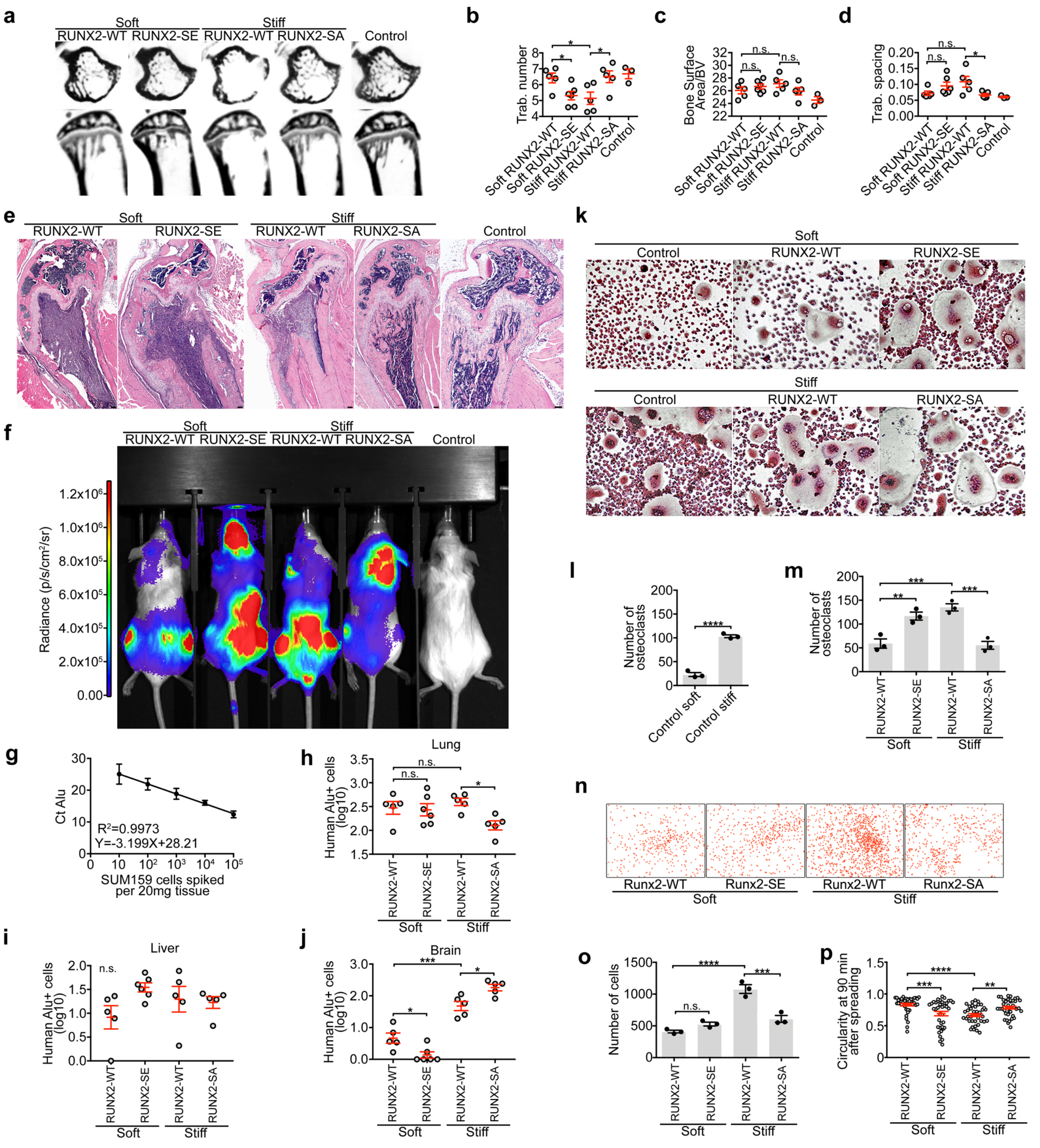


### EXTENDED DATA FIGURE 11

**Extended Data Fig.11. RUNX2-mediated mechanical memory instructs osteolytic bone metastasis. a,** Two-dimensional cross-sections of tibia from mice bearing no cancer cells (control) or SUM159 cells overexpressing RUNX2-WT, RUNX2-SE, or RUNX2-SA, preconditioned for 7 days on soft or stiff hydrogels as indicated, 4 weeks after intracardiac injection. **b-d**, Quantification of trabecular number (**b**), bone surface area/bone volume (**c**), and trabecular spacing (**d**) from mice in (**a**) (*n*: mice; soft RUNX2-WT 5; soft RUNX2-SE 6; stiff RUNX2-WT 5; stiff RUNX2-SA 5; control 3). **e,f,** H&E staining (**e**) and bioluminescence (**f**) in mice from (**a**) noting the strong signal in the skull (second from left) and brain/soft tissue (second from right). **g**, Standard curve from qPCR of human *Alu* using SUM159 cells spiked into 20 mg of mouse tissue, normalized to mouse *actin* ($n = 3$). **h-j**, Quantification of metastases in lung (**h**), liver (**i**), and brain (**j**) from mice in (**a**). **k**, TRAP staining of RAW264.7 cells after 7 days incubation: 4 days with 50 ng/mL RANKL in growth media, and then 3 days with 50% SUM159 conditioned media (CM) + 50% growth media. CM was collected 24 hours after addition to hydrogels with equal SUM159 cell counts in each experimental group. **l, m,** Quantification of (**k**) ($n = 3$ biological replicates with $n = 3$ technical replicates). **n,** Large-stitched images of DAPI-stained (pseudo-colored in orange) SUM159 on synthetic bone matrix after 30 minutes of adhesion challenge. **o,p,** Quantification of cell adhesion (**o**) and circularity (**p**) from (**n**). ($n = 3$ biological replicates with $n = 3$ technical replicates). Data are mean ± s.e.m. *$P < 0.05$; **$P < 0.01$; ***$P < 0.001$; ****$P < 0.0001$, one-way ANOVA with Holm-Sidak's multiple comparisons test (except for **l**: two-tailed unpaired Student's *t*-test). See Source Data for exact *P* values.

#### SUPPLEMENTARY MOVIE LEGENDS

**Supplementary Movie 1.** Time lapse movie of cytoskeletal dynamics of iRFP-Lifeact-expressing SUM159 cells preconditioned on stiff and/or soft hydrogels as indicated in Fig1c. Time is hrs:min:sec. Scale bars = 10 μm.

**Supplementary Movie 2.** Time lapse movie of 3D *in vitro* invasion of SUM159 cells preconditioned on stiff and/or soft hydrogels as indicated in Fig1g. Images are Extended Depth-of-Focus reconstructions of z-series of DIC images. Time is hrs:min:sec. Scale bars = 100 μm.

**Supplementary Movie 3.** Time lapse movie of 3D *in vitro* invasion of SUM159 cells preconditioned on stiff and/or soft hydrogels and FACS-sorted for CellVue proliferation tracker as indicated in Fig3g. Images are Extended Depth-of-Focus reconstructions of z-series of DIC images. Time is hrs:min:sec. Scale bars = 100 μm.

**Supplementary Movie 4.** Time lapse movie of cytoskeletal dynamics of iRFP-Lifeact-expressing SKBR3 and MS-SKBR3.1 cells, preconditioned on stiff and/or soft hydrogels as indicated in Extended Data Fig8a. Time is hrs:min:sec. Scale bars = 10 μm.

**Supplementary Movie 5.** Time lapse movie of 3D *in vitro* invasion of SKBR3 and MS-SKBR3.1 cells, preconditioned on stiff and/or soft hydrogels as indicated in Extended Data Fig8g. Images are Extended Depth-of-Focus reconstructions of z-series of DIC images. Time is hrs:min:sec. Scale bars = 100 μm.

**Supplementary Movie 6.** Time lapse movie of 3D *in vitro* invasion of SUM159 cells overexpressing RUNX2-WT, RUNX2-SE, or RUNX2-SA, preconditioned for 7 days on soft or stiff hydrogels as indicated in Extended Data Fig9a. Images are Extended Depth-of-Focus reconstructions of z-series of DIC images. Time is hrs:min:sec. Scale bars = 100 μm.

**Supplementary Movie 7.** Time lapse movie of SUM159 cells overexpressing RUNX2-WT, RUNX2-SE, or RUNX2-SA, spreading on synthetic bone matrix, preconditioned for 7 days on soft or stiff hydrogels as indicated in Fig4e. Time is hrs:min:sec. Scale bars = 100 μm.
